# Systematic non-natural base-pairs for strand displacement circuits in complex environments

**DOI:** 10.1101/2025.06.28.662084

**Authors:** Tiernan Kennedy, Thomas Mayer, Friedrich C. Simmel, Chris Thachuk

**Affiliations:** Paul G. Allen School of Computer Science & Engineering, University of Washington, Seattle, WA, USA; Physics of Synthetic Biological Systems, Department of Bioscience, TUM School of Natural Sciences, Germany

**Keywords:** Strand displacement (SD), XNA, non-natural bases

## Abstract

Nucleic acid strand displacement (SD) circuits have enabled impressive realizations of molecular computing *in vitro*, but operating them in contention with competing reaction pathways can cause signal leak and circuit failure. Further, operating SD circuits in complex environments, ranging from pools of nucleic acids to living organisms, can exacerbate these challenges. To address these issues, we explore the incorporation of isoC and isoG non-natural bases in single- and multi-step SD cascades and started by studying their promiscuity. We combined the non-natural bases with a mismatch strategy, showing them to be fully compatible. The synthesis we show between strategic placement of non-natural bases and systematic mismatch optimization of SD circuits offers a design space for future circuits that are fast, robust, and biologically compatible. Comparing circuit slowdowns by random pool backgrounds in the presence and absence of non-natural base pairs reveals a significant difference. Using isoC:isoG base pairs in the toehold region shows the greatest effect, with some further positive effects caused by additional non-natural bases in the branch migration domain. Such insights will pave the way for developing SD circuits that operate in complex environments, expanding the applications of molecular computing in a wider range of scenarios.

## Introduction

Molecular computing has emerged as a promising field for information processing at the molecular level, with applications in synthetic biology, diagnostics, and biosensing (2– 4). Strand displacement (SD) circuits, in particular, have enabled impressive realizations of molecular computing *in vitro*, showcasing their potential for complex computations. These circuits, based on the principles of DNA strand displacement, offer a versatile and programmable platform for implementing logical operations and computation. Figure 1 highlights the principles of strand displacement.

**Fig. 1.**
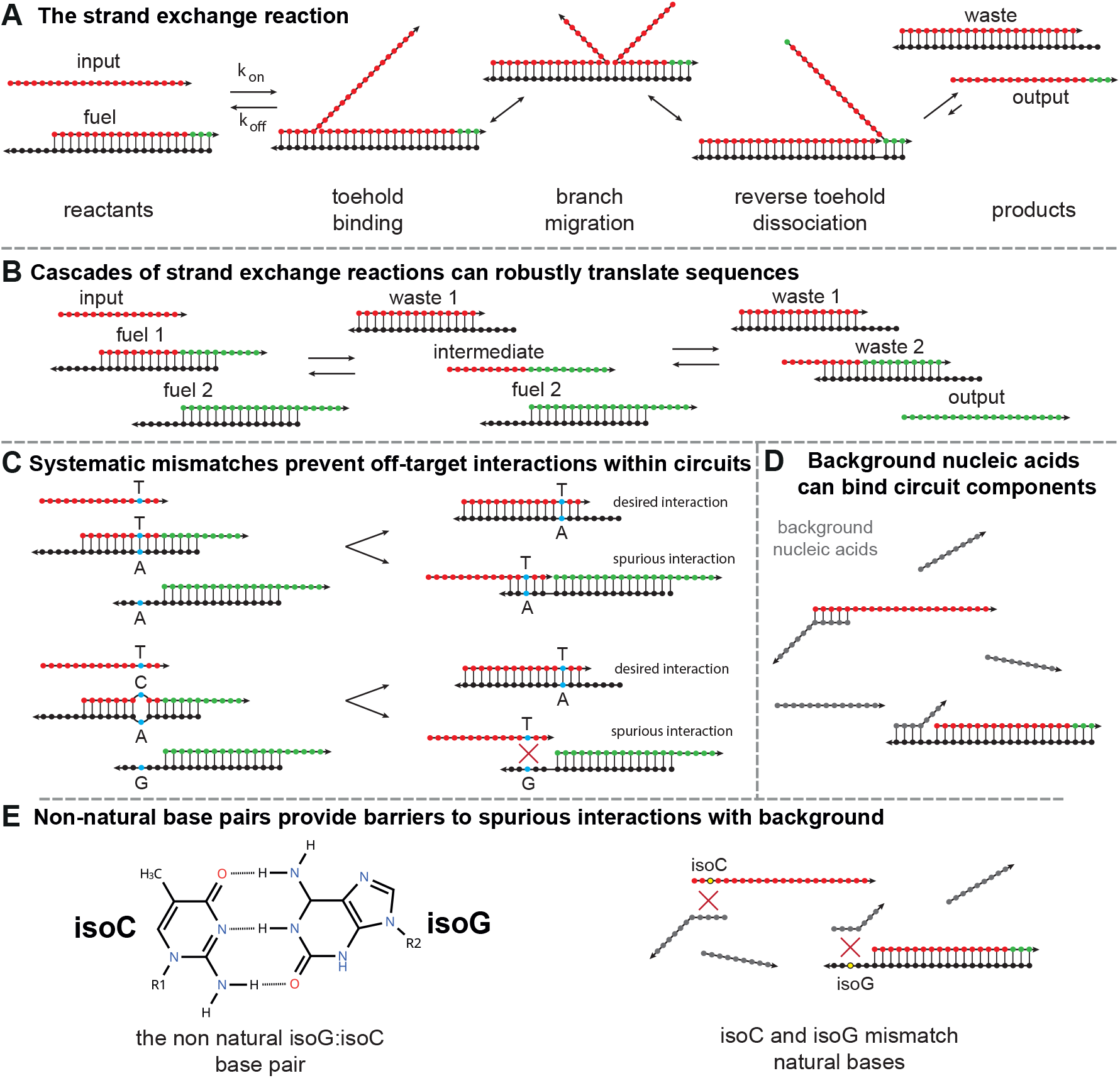
(A) The strand exchange reaction is a versatile tool for engineering chemical reaction networks. The reaction starts when a single-stranded input binds a doublestranded complex (fuel) with a short single-stranded overhang, called the ‘toehold’. After toehold binding, the invading strand (input) can begin to displace the incumbent strand, which initiates branch migration, i.e., a random walk process in which both the invader and incumbent compete for binding to the substrate strand. After the invader is completely bound, the last bases, called reverse toeholds, of the incumbent strand fray to release the output. The ratio of released output, as well as the kinetics, are defined primarily by the different interaction strengths between input and substrate vs. incumbent and substrate. If there is no reverse toehold, strand exchange is typically called strand displacement. (B) Multiple strand displacement or exchange reactions can chain together into a cascade. In a cascade, the incumbent strand displaced from a fuel species further reacts with a downstream fuel. By systematically sharing sequences between fuel complexes, cascades can robustly translate an input sequence to a distinct output (1). (C) The systematic redundancy that imparts robustness to cascades also introduces side reactions between signal strands and fuel complexes, such as that depicted between the input and fuel 2. However, systematic modification with mismatching bases imposes barriers to these side reactions. (D) Even robust cascades that are kinetically optimized with mismatching bases face further kinetic drawbacks when operated in complex biological or environmental samples where they interact with background nucleic acids. These background sequences can bind, e.g., occlude toehold bases, causing operational slowdowns. (E) Xenonucleic acid (XNA) non-natural base pairs, like the isoG:isoC pair, form a complementary base pairing system but substantially mismatch with natural bases. As a result, these bases confer protection against binding by natural nucleic acids. Synthesizing isoG:isoC pairs with systematic mismatch optimization of strand exchange cascades ameliorates slowdowns from competing background nucleic acids by providing significant barriers to binding by any sequence without the complementary XNA.

The simplicity and predictability of SD circuits have contributed to their widespread use and success in various applications (5). Though SD circuits have demonstrated success *in vitro*, operating them in complex environments poses unique challenges (6–8). These environments include pools of random nucleic acids (6), cell extract (9), and living organisms. In these scenarios, SD circuits must contend with myriad factors, such as the presence of competing nucleic acid species, time-varying background compositions, and potential degradation mechanisms (9–14). The complexity and dynamic na-ture of these environments introduce additional constraints and considerations for the design and operation of SD circuits.

The robustness and stability of SD circuits in complex environments are crucial for their practical applications. Traditional SD circuits rely on the Watson-Crick-Franklin base pairs (A:T and G:C) for their operation. These bases can be susceptible to undesired effects and circuit failures when interacting with natural sequences and environmental factors. For instance, unintended interactions may lead to signal leak, false positives, or compromised circuit functionality (15, 16). Additionally, natural degradation mechanisms can degrade circuit components, leading to diminished performance and limited operational lifetimes.(9–14).

To overcome the aforementioned challenges and enhance the robustness of SD circuits in complex environments, researchers have explored incorporating non-natural base pairs. These artificial base pairs offer distinct advantages, such as weakening interactions with natural sequences and slowing down natural circuit degradation mechanisms (9, 10, 14). By incorporating non-natural base pairs, SD circuits can potentially mitigate unwanted interactions and increase their stability and reliability in complex environments.

One such non-natural base pair is isoC:isoG. The isoC:isoG system has been incorporated into both the 8 letter ‘Hachimoji DNA’ (17) alphabet and a larger 12-letter alphabet (18). Hachimoji DNA includes the standard biological bases (A, T, C, G) and two non-natural pairs, P:Z (5-Aza-7-deazapurine : 6-Amino-5-nitropyridin-2-one) and B:S (isoG : isoC in RNA and isoG : 1-MethylC in DNA), allowing for a completely non-natural analogue to DNA.

Incorporation of non-natural bases also has advantages when operating displacement circuits with hundreds of components, which are used for complex computation and pattern recognition in applications like neural networks (19). In such circuits, mostly short toeholds, e.g., of 5 nucleotides (nts), are used to minimize unintended binding to other sequences. However, for such toeholds, the total number of different sequences is only 4^5^ = 1024, and the number of perfectly orthogonal sequences is much lower. By incorporating two new bases, the number increases more than seven-fold to 6^5^ = 7776, and the number of orthogonal sequences also dramatically increases, simplifying the design process. In this study, we investigate the influence of incorporating isoC:isoG pairs into SD components.

This approach is supported by a previous study which showed that building a strand displacement complex with acyclic bases in the toehold reduced binding by natural sequences (20). However this type of XNA in not commercially available, does not interface with enzymatic machinery, and suffers kinetic disadvantages compared to ‘Hachimoji’-style XNA pairs.

We combine the non-natural base incorporation with a general systematic mismatch strategy for fast and robust circuits (16). This strategy, illustrated simply in Figure 2(C) improves kinetics in strand displacement cascades by favoring on-target reactions– which replace mismatches with a matching pair– and disfavoring off-target interactions between circuit components, by ensuring they must form new mismatches. Figure 6 shows an example of a strand displacement cascade systematically optimized with this strategy at multiple sites carefully chosen to maximize kinetic discrimination between on-target and off-target reactions. The strategies proved to be fully compatible, providing additional benefits, beyond those provided by systematic mismatches alone, when operated in complex environments. We present data for single strand displacement reactions and multi-displacement cascades with non-natural base pairs incorporated in the toehold. Furthermore, we show that non-natural bases in the branch migration domain provide some minor benefits, as well. Figure 1 highlights SD interactions, including strand exchange, mismatch strategy, non-natural base incorporation and interaction with background strands. We use three different methods to prepare systems for kinetic analysis, which we present in Figure 2.

**Fig. 2.**
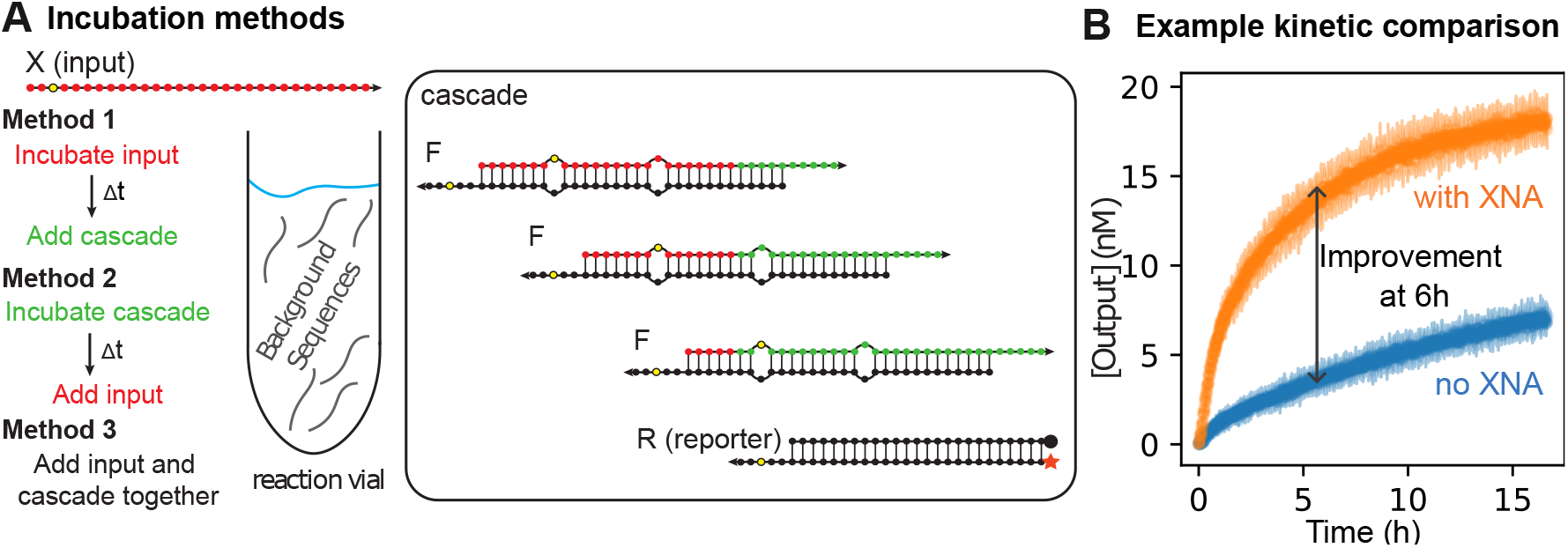
A strand displacement cascade experiment that incorporates non-natural base pairs. (A) Schematic overview of the experiment’s procedure. Either the trigger or cascade and reporter were incubated overnight with background sequences, and the reaction was later triggered by adding the remaining components. Alternatively, no incubation was performed, and all components were added together. (B) Exemplary curves of fluorescence intensity measured over time, normalized to the fluorescence of 20 _nM_triggering in the absence of background for each reporter. In the shown case and throughout this work, the cascade with non-natural base pairs shows increased kinetics in all cases. Comparing the fractional completion in the same conditions with and without the non-natural modification provides a convenient proxy for the extent of kinetic restoration since the controls without background complete before this time-point.

**Fig. 3.**
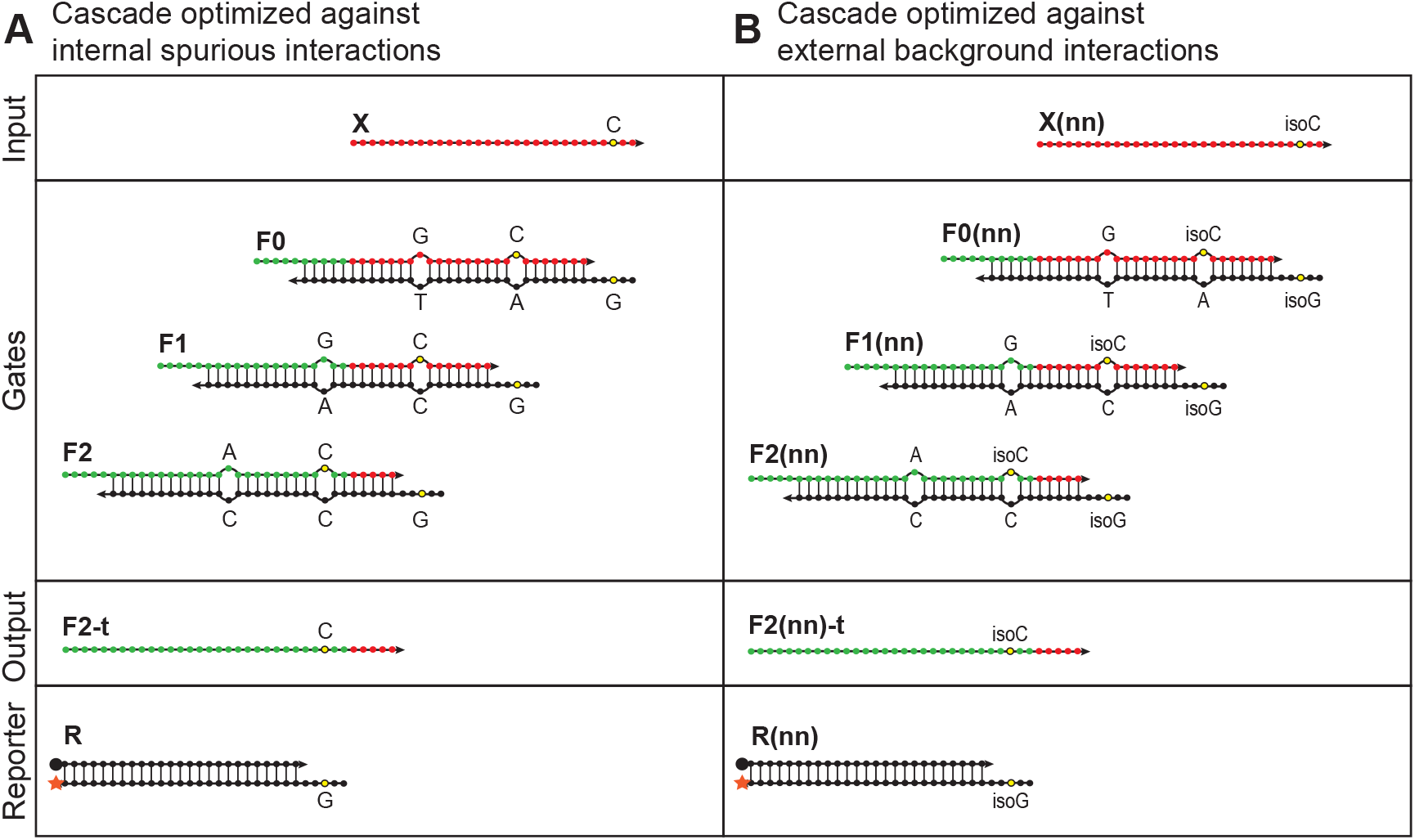
Strategies for achieving robust molecular circuits and robust biological detection. Recent work (16) has shown how to achieve systematic sequence-level optimization of robust SD cascades with internal base pair mismatches. The sites used for systematic mismatch optimization provide a natural choice for introducing non-natural base pairs to optimize circuits simultaneously against (A) internal spurious interactions between components and (B) external spurious interactions with natural nucleic acid background. The bases shown in yellow are modified to be non-natural. A second cascade with the same domain structure but opposite strand directionality and different sequences was also constructed. We present sequences in the supplementary information S4.

Using data from experiments on SD circuits containing non-natural bases in different background environments, we provide insights into altered interactions with background sequences and point out the potential for more robust SD circuits. The kinetic curves in Figure 2(B) show the general trend of faster displacement kinetics for circuits with the non-natural isoC:isoG pair instead of the natural C:G pair in the toehold. Comparing the kinetics of the displacement circuits reveals that background sequences have less influence on the cascade with the non-natural base strategy in all cases.

We designed the different SD cascades with differing toehold strengths and positions (5’ and 3’) to probe the impact on non-natural base incorporation on a broad design space. The 5’ toehold variant is illustrated in Figure 6, and the sequences for both systems are presented in the supplementary information. We compare the kinetics of the SD systems in the presence and absence of pools of random DNA background sequences. Notably, though non-natural bases show promise for engineering bio-orthogonal nucleic acids, in biological conditions these expanded alphabets are not completely orthogonal to natural bases (21, 22).

As part of this work, we investigate the orthogonality of all possible base pairing interactions between natural bases A, T, C, G and the non-natural isoguanine and isocytosine as demonstrated by their influence on SD kinetics when substituted in the central position of a 5 nt toehold. For better comparison, we perform the same experiments with the natural C and G interacting with all possible natural bases. Our results show that even a single non-natural base in each toehold of a SD system significantly reduces the negative impact of random background sequences. Our characterization of the relative orthogonality of different non-natural:natural pairs and additional benefits caused by non-natural bases in the branch migration domain provides a solid foundation for engineering robust strand displacement circuits with optimized orthogonality to pools of nucleic acid background. Furthermore, the synthesis we show between strategic placement of non-natural bases and systematic mismatch optimization of robust strand displacement circuits offers a design space for future circuits that are fast, robust, and biologically compatible.

## Methods

We were interested in having different toehold placements, which lead to each lab using different cascade designs and some differences in the methods used. One lab conducted all experiments on the 3’ toehold cascades, while the other conducted them on the 5’ toehold cascades. The versions (3’ or 5’) are specified in the following discussion.

### Material preparation

Where applicable, existing designs from (6) for the 5’ toehold and (16) for the 3’ toehold cascades were used. The reporter for the 5’ version had an incumbent with a 5’ Iowa Black^®^ RQ and a substrate with a 3’ ROX (NHS Ester) fluorophore. The 3’ version used a 3’Black Hole Quencher modification on the incumbent strand and a 5’ ATTO565 modification on the substrate strand.

All main SD-circuit components were ordered either PAGE purified or HPLC purified and lab ready at 100 µM except for the natural variants for the multi-isoC and long overhang systems, which were ordered unpurified and were later PAGE purified by hand. Sequences with random base compositions at certain lengths (N25 and N50) were ordered as dried DNA and renatured in 1x TE buffer with 100 mM NaCl, 12.5 mM MgCl_2_ and 0.01% Tween 20 for the 5’ toehold system experiments and 1x TE buffer with 12.5 mM MgCl_2_ and 0.01% Tween 20 otherwise. The reporters were annealed with 1.2X excess top strand and used without further purification. All upstream complexes were annealed with 1.2X excess top strand and purified by native polyacrylamide gel electrophoresis to recover the band corresponding to the duplex.

### Experiment conditions

The reporter consisting of substrate and incumbent was always used at a final concentration of 40 nM, the gates at 30 nM, and the trigger at 20 nM. For the 5’ version, fluorescence measurements were performed at 29^*°*^C using either a FLUOstar^®^ or CLARIOstar^®^ plate reader by BMG LABTECH and Corning^®^ low volume, non-binding plates. The filters were chosen to be 584 nm for excitation and 640 nm for emission.

For the 3’ version, reaction components and solvent were combined using a Labcyte Echo525 acoustic liquid handling robot into a Corning 384-well black clear bottom microplate. All fluorescence experiments for this version were conducted using BioTek Synergy H1 fluorescence or BioTek Cytation 5 plate readers set to 25 ^*°*^C, with excitation and emission wavelengths optimized for ATTO565.

For all measurements containing random pool strands, we used three different incubation methods: none, trigger, or cascade. For trigger and cascade incubation, the system equilibrated overnight.

### Data normalization

To account for the different conditions and material used in the experiments, we always compared the measurements within experiment subsets that had the same conditions. To improve readability, we normalized the intensity data to match the signal for the reporter or cascade triggered with 20nM input in the absence of background. The concentration of the input strand stocks were determined through triplicate absorbance measurements taken at 260nm using a Thermo Scientific™ NanoDrop™ One Microvolume UV-Vis Spectrophotometer. Since the extinction coefficients were not available for the isoC and isoG bases, we approximated the extinction coefficient of these non-natural strands as identical to their natural analogue because the sequence difference contributes only a small percentage to the overall extinction coefficient for the strand.

For each measurement, we subtracted the intensity value for the un-triggered reporter (when all fluorophores should be quenched) to set the zero value. We then normalized fluorescence value such that the difference in fluorescence between the reporter/cascade triggered at 20 nM in the absence of background and the baseline un-triggered reporter fluorescence equals 20. The fractional completion was computed at 6 hours of reaction time.

For the 5’ toehold system, after purification the concentrations of the gates in the cascade were estimated using their extinction coefficients values and absorbance measurements with an Implen Nanophotometer N50. For the 3’ system, each gate’s (including the reporter) stock concentration was determined by measuring the fluorescence of the fully triggered gate with the downstream gates in excess. The fluorescence was then converted to a concentration value using a calibration curve of the reporter, kept at the same concentrations as the used in the excess triggering.

### Data modeling

All models were fit to data after normalization to the 20nM control triggering. Errors for the second-order rate constants were estimated using the jackknife method. To achieve this, each possible pair of duplicates for each of the three replicate curves for each experiment condition were averaged together (the set of duplicates achieved by leaving out one replicate at a time). The following ODE model represents a single irreversible reaction, where I is the input, F is the fuel, W is the waste, and O is the output:

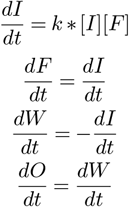

This model was then fit to the first six hours of each experiment’s output curve using the Levenberg-Marquardt Algorithm for non-linear least squares and the experiment’s concentrations as initial conditions. The rate constant was reported as the average of each of the three rate constants from the sets of duplicates, and the error was reported as their standard deviation. As a comparison, the first 15 points of each normalized output curve were fit to a linear model using linear least squares (see supplementary information). The initial rate was reported as the average of the slope fit to each possible subset of experimental duplicates, and the uncertainty was reported as the standard deviation of this set.

## Results

### Mismatches of natural and non-natural toeholds

We conducted displacement kinetic experiments on all possible combinations of the non-natural isoC and isoG bases with all natural bases, as well as for the natural counterparts C and G. For comparison, we added the matching isoC:isoG kinetics, which proved to be the fastest, followed by the combination of the natural C:G pair. Figure 4(A) shows the kinetic curves for all combinations with the non-natural bases, and completion levels for all combinations including the natural mismatches are shown in 4(B).

**Fig. 4.**
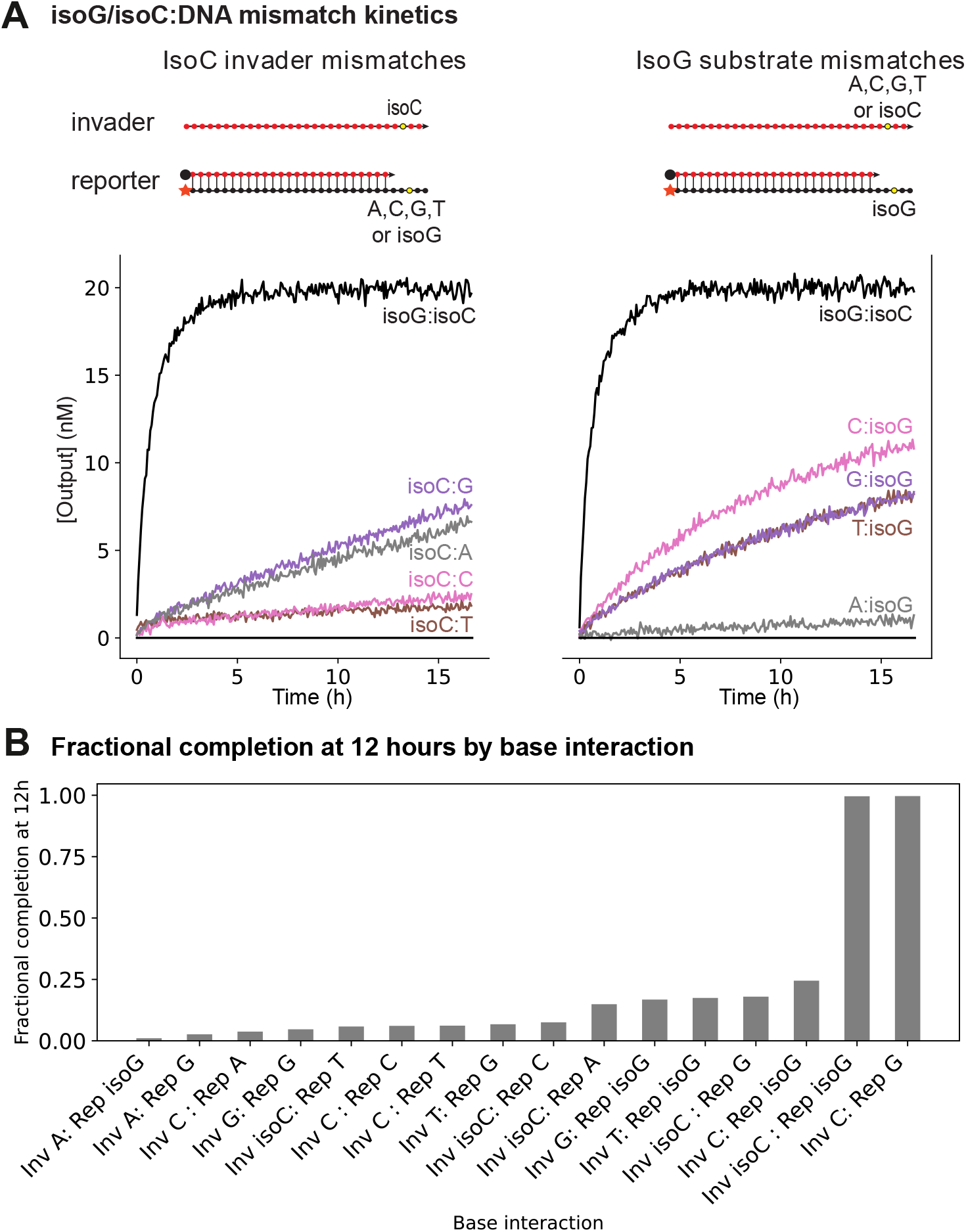
Mismatches with isoC or isoG show increased promiscuity compared to natural-only mismatches. (A) Kinetic curves measured for the 40nM reporter and 20nM trigger with various matching and mismatching bases in the center of the toeholds. (left) Data for the isoC base in the invading strand mismatching to each of the standard A,T,C,G bases or matching the isoG base in the center of the reporter toehold. (right) Analogous data for an isoG base in the center of the reporter toehold. (B) The promiscuity of each mismatch of isoG and isoC, assessed by the completion level at twelve hours in the experiments from (A) compared to the analogous data for each possible mismatch of the natural C and G bases in the center of the toehold. We observed significantly increased promiscuity of isoC and isoG compared to natural C and G.

In this case, observable triggering kinetics are still possible since only the middle of the toehold is not matching, but the remaining 4 bases are still complementary. However, for the used, weak toehold (see SI Table S4 for all sequences), one mismatch is sufficient to prevent initial, stable toehold binding. For the natural C base, only the matching G base significantly increases fluorescence intensity for the measured timescale. For isoC, the combination with the natural G base, as well as with the natural A base, increases the fluorescence signal. The fact that G and A showed some interaction with the isoC base could be because they are both purines and therefore share some chemical characteristics.

For the combinations of G with the natural bases, the G:T and G:G mismatches show some interaction, although the latter is barely visible. For isoG, noteworthy interactions occur for several combinations. The strongest is for isoG:C, which shows slightly faster kinetics compared to isoC:G, which is the strongest interaction for the aforementioned case. This is consistent with the findings by Roberts et al. (23).

For isoG:T and isoG:G, the fluorescence increase is equally pronounced. The increased promiscuity of isoG compared to isoC is consistent with the increased number of theoretical low-energy tautomers capable of binding the natural bases. This is qualitatively consistent with previous melting studies of duplexes that incorporate isoguanine and 5-Metyl isocytosine bases (24). However, we observed a different order for the favorability of isoguanine mismatches compared to the duplexes used in that prior study. Our use of different sequences around the non-natural base sites, compared to Bande et al, may indicate significant differences in energetics depending on the nearest neighbor context.

### Influence of random pools on single-step displacement kinetics

To gain insights into the effect of non-natural base pairs on the kinetics of SD circuits with random pool strands, we measured different combinations, varying the length (N25 and N50) and concentration (10 and 25 µM) of the random pool strands in combination with a single strand displacement reaction. For these experiments, we used the strands of the last strand displacement in the cascade shown in 3 and the different incubation methods described in the methods section, since one component in a sensing application is in the cell or complex sample and is therefore already equilibrated with the surrounding. We also compared the incubated case to a non-incubated variant to investigate the degree to which the background binding equilibration influences the bulk kinetics.

We extracted the data for the completion levels after 6 hours for the system with and without the systematic incorporation of non-natural bases, as shown in Figure 2(B). We then compared the percent of the completion level the systems had reached at that time, as we show in Figure 5(B) and (C). We observe that the displacement circuits with the non-natural bases show an increased completion level for all cases.

**Fig. 5.**
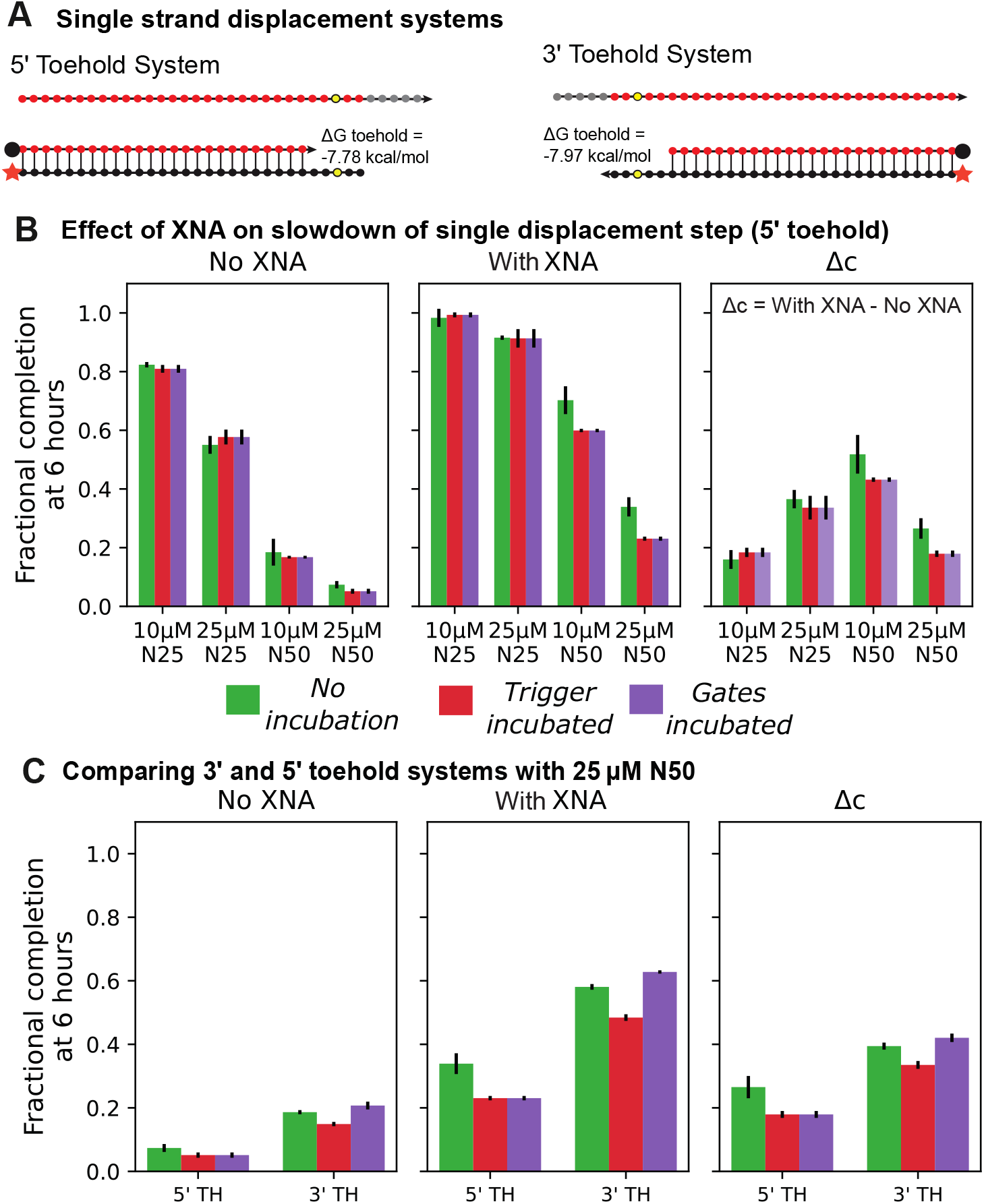
Non-natural base incorporation improves performance of a single strand displacement in random pools of nucleic acids. (A) Sequence depiction of the two different single displacements used for this set of experiments, including their toehold strength in the case with only natural bases. (B) Comparison of the fractional completion at 6 hours for the 5’ toehold version and different background lengths and concentrations, as described in the Methods section. For easier comparison, the rightmost graph shows the increase in fractional completion by subtracting the completion without non-natural bases from the completion with them. (C) Comparison of the 5’ and 3’ toehold versions for one example background case (N50 at 25 µ_M_). The 3’ toehold version reaches a higher value after 6 hours with and without non-natural bases, however both versions notably benefit from incorporating the isoC:isoG pair.

**Fig. 6.**
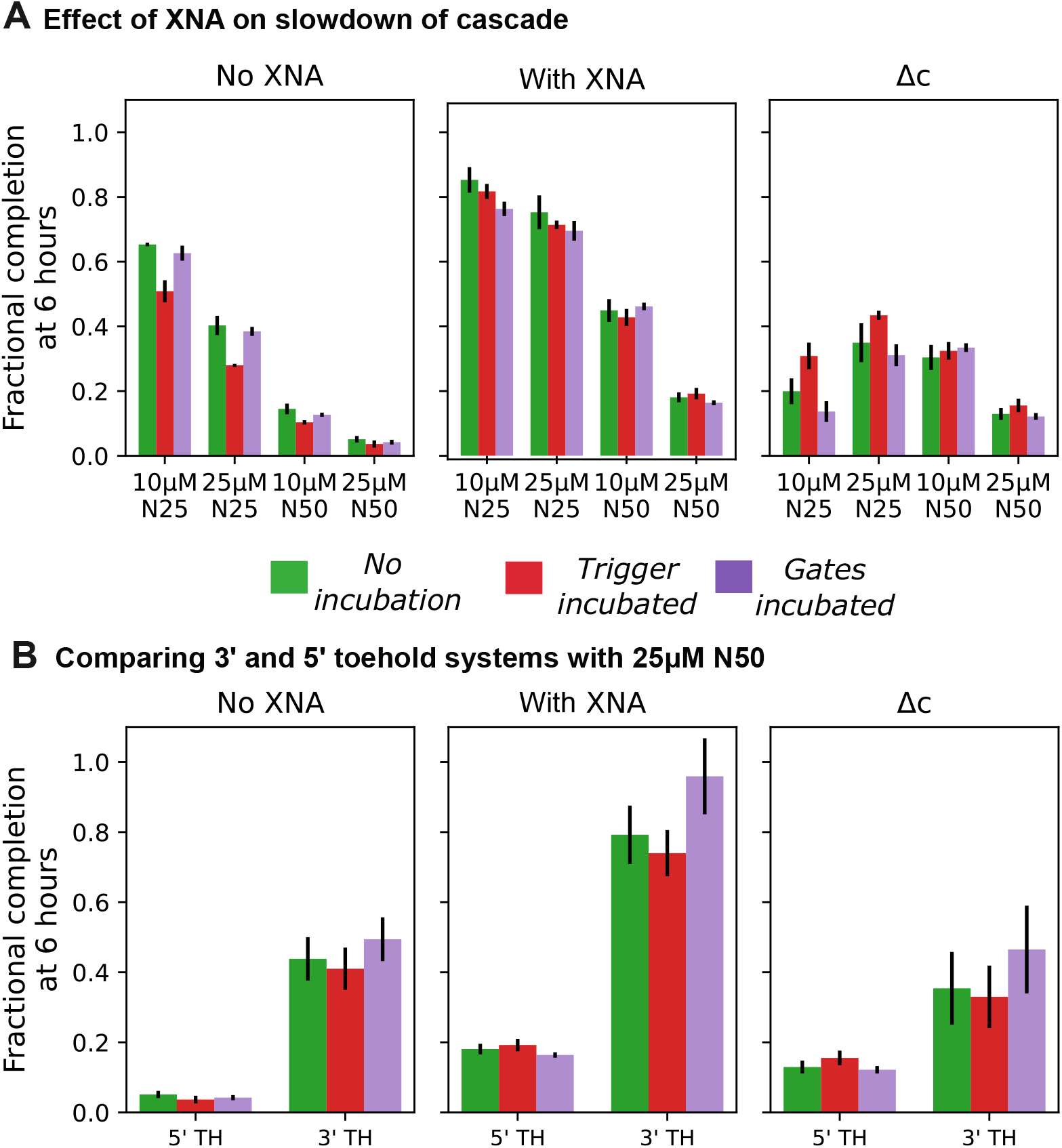
IsoC/isoG incorporation improves the performance of larger SD cascades in the presence of random pools of nucleic acids. (A) Comparison of the fractional completion at 6 hours for the 5’ toehold version and different background lengths and concentrations, as described in the Methods section. For easier comparison, the rightmost graph shows the increase in fractional completion by subtracting the completion without non-natural bases from the completion with them. (B) Comparison of the 5’ and 3’ toehold versions for one example background case (N50 at 25 µM). As already seen for the single strand displacement reaction, the 3’ toehold version reaches a higher value after 6 hours, which is even more pronounced for the multi-step cascade. However, both cascades significantly benefit from incorporating the isoC:isoG base pair in the toehold.

Kinetics with and without background were significantly faster for the 3’ vs the 5’ systems, despite the similar toehold binding energies for both reporters. This could occur because strands in phosphoramidite synthesis are created 3’ to 5’; therefore, truncations are most likely at the 5’ end, which could cause the 5’ toehold to behave as if it were shorter than predicted. The fact that incubating the reporter for the 3’ system led to the larger completion increase than the trigger incubation, which is not the case for the 5’ toehold system, also supports the notion of a qualitative difference between the toeholds of the two systems.

Regardless of the cause of this kinetic difference, the system with the faster kinetics also showed a larger increase in completion from incorporating the non-standard bases. In general, individual incubation kinetics that were faster without background showed a larger increase in completion at 6 hours, likely due to the exponential nature of a kinetic trace contrasted with a fixed time-point completion measurement.

### Influence of random pools on cascade displacement ki-netics

We have established that non-standard bases offer a significant improvement in the kinetics of strand displacement in the presence of a random DNA background. It is natural to question whether this holds true for larger displacement systems with multiple displacement steps. In this set of experiments, we investigate different techniques for adding/incubating strands with the random pool.

In cellular sensing, receptors sample the environment and signal is propagated only when a ‘sufficiently intense’ stimulus is encountered. In this case, the inputs are available on short timescales, reducing their likelihood of encountering stable off-target binding partners. The same was also found in synthetic studies of random background in SD circuits (6), where mixing DNA signals with random pool backgrounds immediately before the reaction commences caused less slowdown than circuits with inputs incubated with the background. Thus, it is important that the strand or complex existing on the longest timescale has no strong interaction with the background.

We present evidence below that incorporating non-natural bases into the sequences reduces interference from random natural nucleic acid pools in all tested cases, i.e., no incubation, incubation of the trigger or incubation of the cascade and reporter. Thus, if we incorporate non-natural bases strategically into toehold sequences, we can reduce the affinity to background sequences.

First, we did not observe any leak, or output in the absence of trigger, for either cascade and any of the concentrations of random background tested in these experiments. (Example kinetic curves showing the absence of leak are included in the supplementary data Figure S1). Second, as in the single strand displacement reaction, we found that the circuit containing isoC:isoG was significantly faster than the natural counterpart. (We include fit rate constants for each complex in both cascades for the 3’ version in the supplementary data Figure S9.) Third, despite the differences in kinetics, we observe that both cascades completed their reaction by 6 hours, so the difference in completion between cases with random background serves as an interesting proxy for the effect of background on the cascade. Fourth, we found that incorporating isoG and isoC bases systematically into cascades provided similar kinetic benefits in the presence of random pools of DNA background compared to the single strand displacement reaction. As shown in Figure 6, we observed a comparable increase in completion for the multi-step 5’ toehold cascade as we did for the single displacement. The data for the 3’ toehold version shows the positive effect on displacement kinetics, as well.

To provide another way to compare the kinetics than by looking at the fractional completion, we also used another fitting method and compared the ratios. We fit second-order rate constants (bimolecular fits) to the fluorescence data. Calculating the rate constant ratio by dividing the rate for the case with non-natural bases by the rate for the case without non-natural bases gives equal or higher values (meaning less influence by the background strands) than the ratio without background for all cases. These values also generally increase with increasing background length and concentration. Figure 7 shows these results. We also extended this analysis using a linear fit to the first 15 minutes of the data to estimate the initial rate of reaction (see supplementary information).

**Fig. 7.**
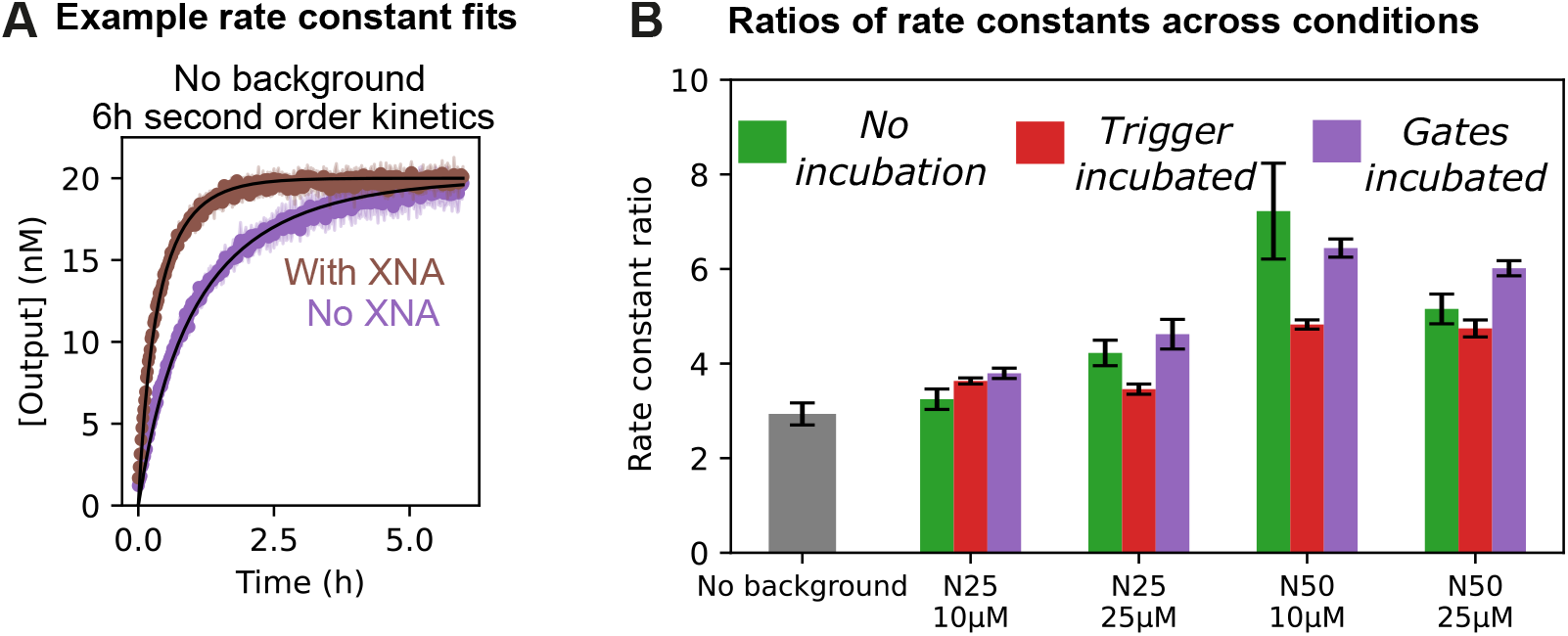
Alternative analysis of the kinetic effect caused by incorporation of the non-natural isoC:isoG base pair. (A) Example of a fitting a second-order rate constant (bimolecular fit) to the collected fluorescence data over the time period of 6 hours. (B) Ratios of the rate constants extracted from the bimolecular fits. The ratio is calculated by dividing the rate for the case with non-natural bases by the rate for the case without non-natural bases. The leftmost bar shows the ratio without the influence of background strands. All other ratios for the three different incubation methods show equal or better performance, meaning less influence by the background strand for the cases with non-natural bases.

The fact that the fold-improvement in the second-order rate constants exceeded the control value without background suggests that the kinetic impact of the XNA goes beyond providing an increased rate constant for the reaction without background. The qualitative differences in the trends for the second-order rate constants and the initial kinetics alone suggest a complicated mechanism for ameliorating reaction kinetics by the non-natural bases that affects a more complicated reaction network than a single bimolecular reaction– including the additional equilibrium of background binding, for example.

### Influence of multiple non-standard bases on kinetic slowdown

We have shown that incorporating non-natural bases in the toehold region is beneficial for SD kinetics. We expanded our investigation by adding non-natural bases in the branch migration domain of signal strands.

To test the impact on kinetics, we designed a reporter with two mismatches and either an A or isoG base in the toehold region; this enabled us to use different triggers with 0, 1 or 2 non-natural bases on the same reporter without significantly changing the free energy of reaction when introducing the second non-natural base in the mismatch position (since no new base pair is formed when substituting one mismatch for another). Using this technique, we could separate the influence caused by the stronger toehold binding and less interaction with the background from the interaction of the branch migration domain with the background. Figure 8 shows the results.

**Fig. 8.**
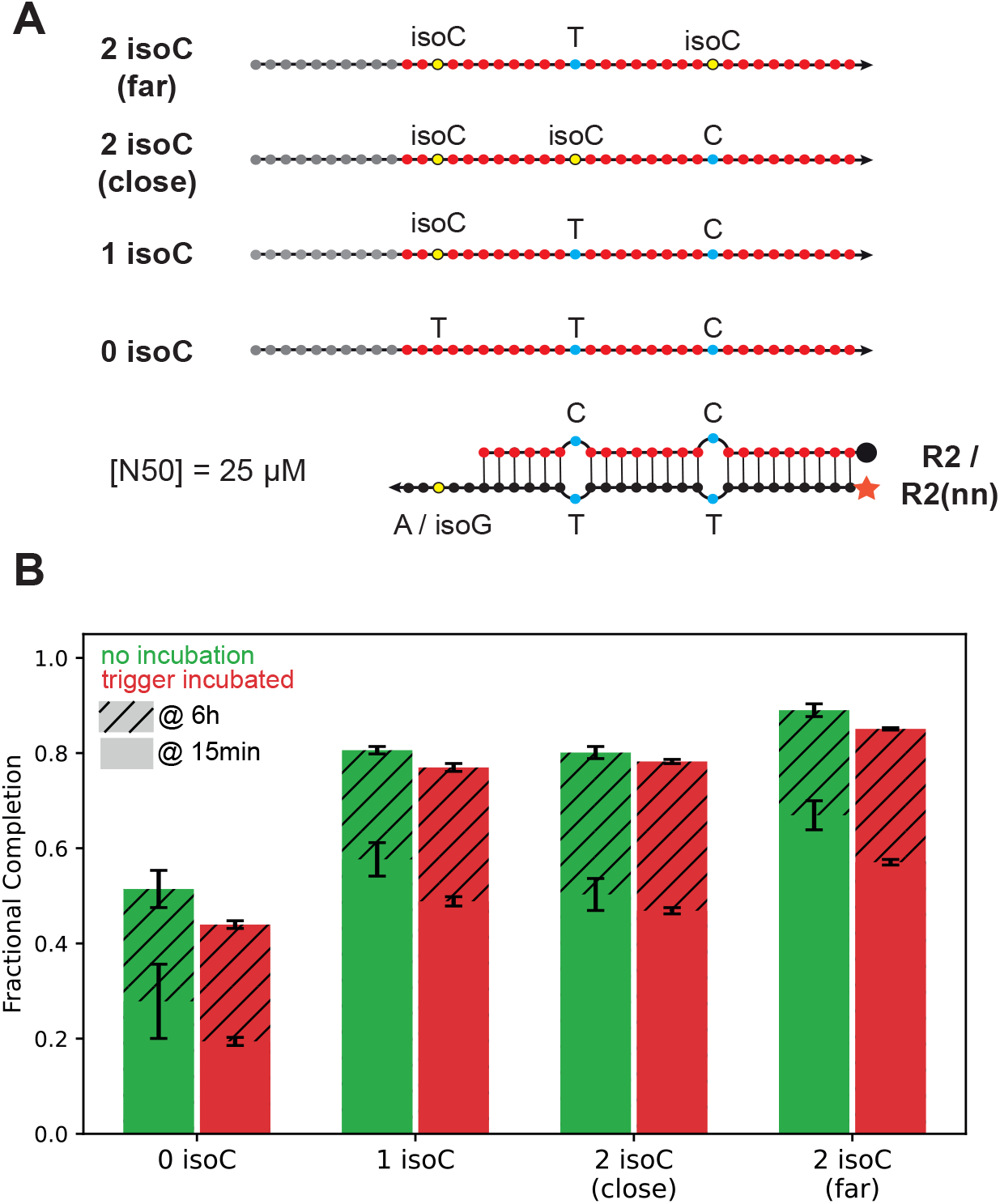
Incorporating additional non-natural bases outside the toehold region of the invader shows only minor kinetic improvements in the presence of random pools of nucleic acids. (A) Sequences used for this set of experiments. The reporter has two mismatches and either an A or isoG base in the toehold region. The mismatches where chosen to avoid distorting the results by providing additional stability for binding one of the invader versions. (B) The fractional completion after 15 minutes and 6 hours for all versions with N50 background strands at 25 µM. Introducing the isoC:isoG base pair in the toehold has major benefits for SD kinetics, whereas the second non-natural base shows only a minor kinetic improvement when placed distal (far) from the toehold and no effect for proximal (close) placement.

Unsurprisingly, introducing the non-natural base pair in the toehold region leads to a significant speedup, but including the second isoC is beneficial only for the case where the isoC is put further away (distal) from the toehold; no difference is visible for the proximal case. We hypothesize that this positional discrepancy might be due to the sequence composition at the mutation sire. For example, the distal isoC disrupts a GG stretch, but the proximal does not. Alternately, inserting the proximal isoC creates the same trinucleotide sequence as in the region over the toehold, which could increase the extent of toehold occlusion (especially with the strong isoG:isoC bond) and slow kinetics. In cases where the second isoC did improve completion, the incremental increase in completion was smaller than that of adding the first isoC over the toehold. Finally, we investigated whether a non-natural base in the overhang of a gate is beneficial for displacement kinetics. Since the cascades we investigated use short 5 nucleotide overhangs that would have a similar binding stability with background as a toehold, we sought to determine whether (1) a longer overhang might lead to a more burdensome interaction with background, and (2) inclusion of an isoC base would be beneficial.

We used the same reporter as described above to ensure differences were not caused by additional stability. The results in Figure 9 show only a very slight improvement in reaction kinetics. This high completion level in the control case and the minimal increase upon the addition of the isoC suggest that overhang sequestration does not pose a rate-limiting step. Combining the minimal further increase from the additional isoC in the signal strands with this result for the overhang indicates that increasing incorporation of non-natural bases yields diminishing returns despite a constant cost per base.

**Fig. 9.**
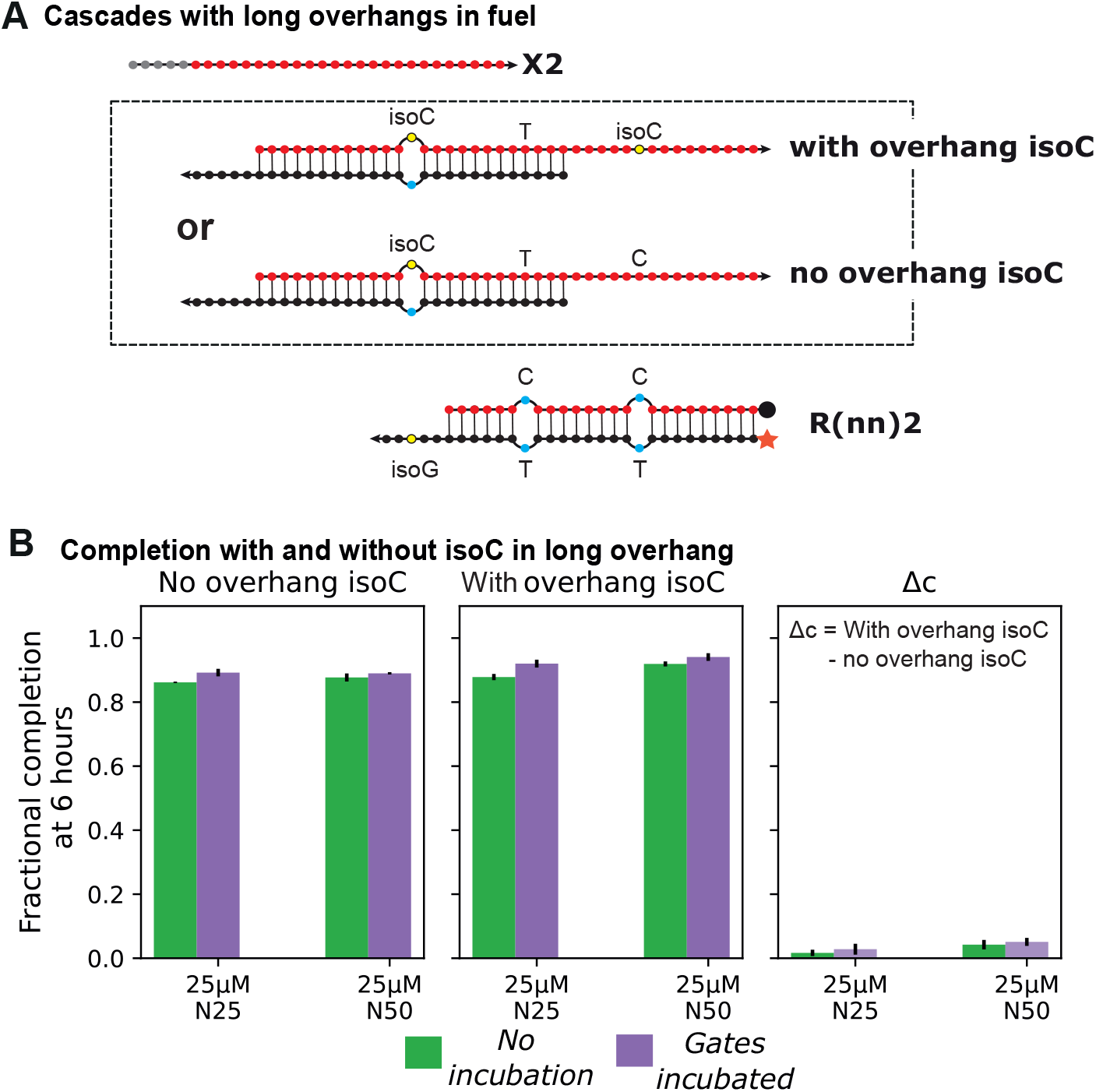
Incorporation of additional non-natural bases in overhangs of strand displacement fuel complexes. (A) Sequences used for this set of experiments. The reporter has two mismatches, which were chosen to avoid distorting the results by providing additional stability for binding one of the invader versions. (B) The fractional completion after 6 hours for the two versions with two different background lengths (N25 and N50) at 25 µM. Comparing the fractional completion for the cases with and without the non-natural base (rightmost graph) shows a negligible difference for N25 and only minor improvement for N50. This suggest that modification of the overhang is not rate limiting compared to modifying the critical toehold region.

## Discussion

Much recent work in the field has kinetically and thermo-dynamically optimized SD circuit components against spurious reactions (1, 16). However, this work does not address how to make components robust against spurious interactions with the background environment. As illustrated in Figure 3, our strategy complements these existing internal optimization techniques to provide additional external optimization.

If starting from a translator or other robust enthalpy neutral design (1) with spurious invasion pathways optimized via systematic mismatches (16), it is straightforward to design isoC/isoG bases into every toehold in the circuit, which we have shown to reduce kinetic slowdown caused by natural sequence backgrounds. Additionally, the strategic sequence positions used for sequence-based optimization of spurious invasion pathways provide a natural choice as the sites for systematically incorporating isoC:isoG pairs at additional positions to promote orthogonality to the background. Together, these strategies might lead to the design of circuits with significantly reduced spurious reactions, both internally to the circuit and externally to the matrix.

Additionally, we demonstrated the effectiveness of our strategy whether the circuit input or the circuit itself is incubated with the background for a long period of time. This is promising for engineering and bio-sensing applications. For example, a designer may want to introduce information encoded as a DNA signal into a complicated environment for a long period of time with the intent of later adding a detection circuit to retrieve the information, e.g., for data storage or tagging applications. Or, a designer might engineer a biosensor that must wait in a complicated background for a long period of time for the correct signal to appear. Our results suggest that strategic incorporation of non-natural bases can improve performance, imparting increased orthogonality to natural nucleic acids in the background for both the input to the circuit and the circuit components themselves.

We also tested whether incorporating additional non-natural bases outside the toehold region adds to the positive kinetic effect. Our results show a slight kinetic benefit that is minor compared to the speedup caused by the non-natural base pair in the toehold region.

While we chose to investigate the isoG:isoG pair because it is widely commercially available, it is more expensive than a natural base pair. Since we found that increasing non-natural bases throughout a design yield diminishing returns, those needing to economize when incorporating non-natural base pairs in their design should consider using one base pair per toehold or placing any further non-natural residues in or around the toehold.

The used random pool concentration in our system might seem high initially, but it is more meaningful in the context of cells. The volume of a cell is much smaller than volume of a bulk solution, leading to much larger effective concentrations of components for the same number of molecules; further, an altered random strand background composition can lead to a more versatile distribution of displacement kinetics, including much slower displacements (25). For example, assuming a compartment with a volume of 1 µm^3^ = 1×10^−15^ L, which is about the size of an E.coli bacterium, a single DNA/RNA molecule (A) would result in a concentration of

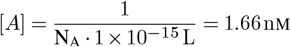

with N_A_ = 6.022×10^23^ mol^−1^ being the Avogadro-constant. This means that only 10,000 nucleic acid molecules would lead to a concentration of 16.6 µM, which has the same order of magnitude as our experiments. Considering that just the number of mRNAs (a small component of the total RNA) in one E.coli cell is already around 1,380 (26), our experimentally tested concentration range seems to be reasonable.

Our investigation of different cascades with different toehold strengths, toehold positions, mismatch substitutions, and altered concentrations has shown a clear common behavior. Slowdown of the displacement reaction was observed in all cases, but introducing the non-natural bases had a positive effect on the kinetics.

However, one additional observation showed surprising deviations from previous work. Incubation of the reporter with the random pool showed almost no influence for the long toehold used in (6) but a very pronounced influence in the cascade system with a weaker toehold of only 5 nucleotides. We hypothesize that this deviation could be caused by a combination of multiple factors. First, occlusion of only one or two bases of the short toehold significantly reduces the initial binding strength, whereas for long toeholds (exceeding 10 bases), one or two occluded bases are not as influential because binding kinetics level at a toehold length of approximately 7 bases, depending on the GC content. Second, finding a sequence stretch of up to 5 complementary bases in the random pool strands is much more likely than finding a subset of, e.g., 10, perfectly complementary bases for a longer toehold. Although binding of 5 bases should not enable a very stable interaction, a pre-incubated system with a large excess of the background could still lead to a favorable equilibrium for background binding.

We characterized the implementation of non-natural base pairs in the toehold region of a DNA SD complex and additional non-natural bases in the branch migration region, including an experimental demonstration of the interplay of non-natural bases with random pools of natural nucleic acids. Doing so provided significant insights into the design and operation of SD circuits in complex environments. We found that incorporating isoC bases into the signal sequence can reduce sequestration of the sequence, leading to faster SD ki-netics even when incubated overnight with a random pool of natural sequences. In the absence of a random pool, reaction kinetics were found to be quickest with the non-natural isoC:isoG base pair and the conventional C:G pair, supporting prior studies of the base pairs’ thermodynamics (27, 28). A critical observation pertains to the tautomeric nature of isoG, which shows some interaction with natural bases, indicating that isoG does not provide perfect orthogonality. Degradation resistance of strand displacement circuits constructed from Hachimoji or other non-natural bases remains an area of future work. Previous work suggests that capping strands with non-natural nucleotides may provide exonuclease resistance; however, Kawabe et al. were able to successfully remove each of 8 non-natural bases with exonucleases as part of their sequencing process (18), so a synthesis of our strategy with L-DNA caps (10) known to provide exonu-clease resistance might provide a promising avenue forward. Non-natural nucleic acids with nonspecific nuclease resistance might be achieved through modified backbones or non-natural bases with significant differences in minor groove electron density (17). Furthermore, even promiscuous endonucleases are known to have natural sequences with reduced endonuclease activity (29); therefore, strategic evolution experiments with non-natural bases may increase the set of known low endonuclease activity sequences or even identify some with lower activity than their natural counterparts. Tools for simulating DNA and RNA thermodynamics, like NUPACK (30), model each base in the alphabet using a collection of nearest-neighbor pair thermodynamic parameters. Further experimental characterization can establish these nearest-neighbor parameters for Hachimoji DNA, L-DNA, and other alternative base pairs. Tools like oxDNA (31)might include non-natural bases like the isoC:isoG pairs as coarse-grained fields in molecular dynamics simulations of DNA.

In conclusion, the strategic and systematic incorporation of non-natural nucleic acids adds to the toolbox for designing and implementing SD circuits. We present promising experimental data in this direction. However, the detailed implications and possible challenges in larger experimental systems require thorough computational and experimental study.

## ACKNOWLEDGEMENTS

We are grateful to Tracy Mallette and Zoe Derauf for their valuable input and suggestions throughout this project.

## Supplementary Note 1: Exemplary Leak

**Table S1.** Leak (output in the absence of input) in the for the 3’ toehold system’s reporters over a period of 10 hours. To differentiate leak resulting from the spurious interaction between gates and fluorescence from transient separation of the fluorophore and quencher on the reporter by background, the signal increase for each reporter in the presence of background was subtracted from the signal for the cascade in the presence of background.

| Reporter type | Background condition | Completion increase over 10 hours |
| --- | --- | --- |
| No XNA | No background | 0.71% |
| No XNA | 25μM N50 | 0.99% |
| With XNA | No background | 1.01% |
| With XNA | 25μM N50 | 0.64% |

**Fig. S1.**
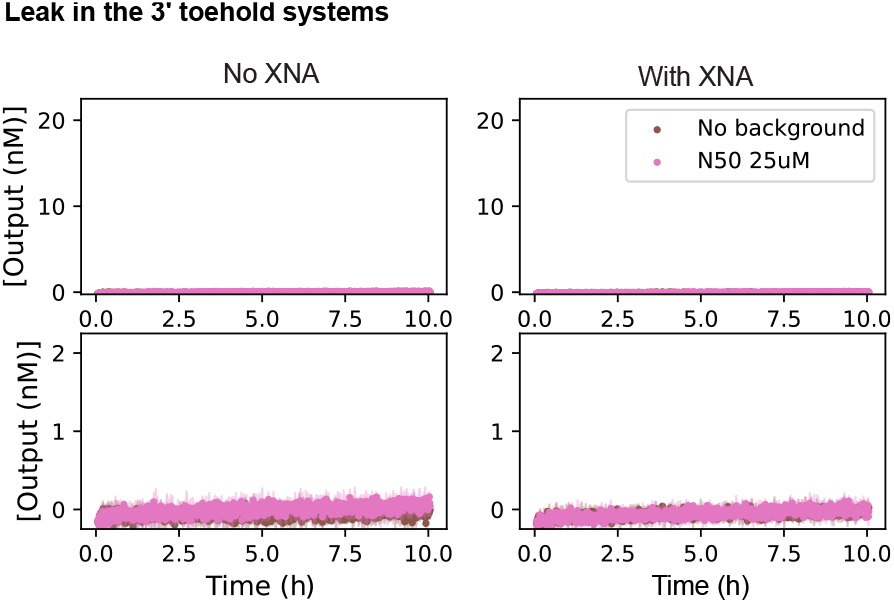
Leak (output in the absence of input) in the for the 3’ toehold cascade over a period of 10 hours. Data for the cascade without XNA is shown in the left column, and the variant with XNA on the right. The pink curves represents leak in the presence of N50 25*µ*M while the brown conditions is in the absence of background. Each row of the figure shows the same data, but the scale of the y-axis in the lower row is zoomed by a factor of ten. Values for quantifying the leak in each case is presented in Table S1. When normalizing the R.F.U. values the reporter, baseline fluorescence without background was used for the cascade without background, and the reporter fluorescence with background was used for the cascade with background. This was done to decouple the increase in reporter auto-fluorescence with background from measuring leak resulting from the spurious interaction between gates.

**Fig. S2.**
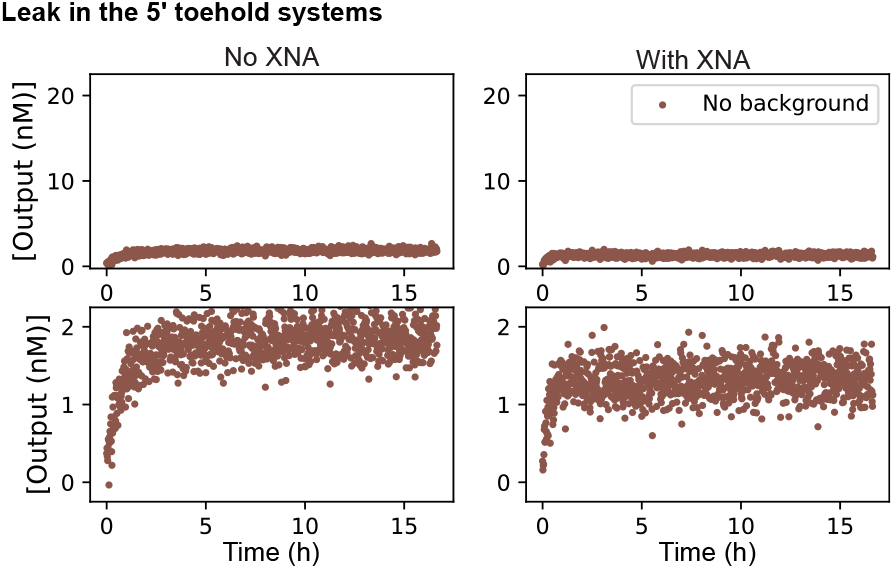
Exemplary leak (output in the absence of input) in the for the 5’ toehold cascade in the absence of background. Data for the cascade without XNA is shown in the left column, and the variant with XNA on the right. Each row of the figure shows the same data, but the scale of the y-axis in the lower row is zoomed by a factor of ten. Both systems show some initial fluorescence increase, which can result from slight inaccuracies in stoichiometry when annealing gates, however after plateauing the curves show a negligible further increase in signal. The inclusion of XNA in the circuit did not increase the initial fluorescence change or long-term leak.

## Supplementary Note 2: Fitting of kinetic constants

**Fig. S3.**
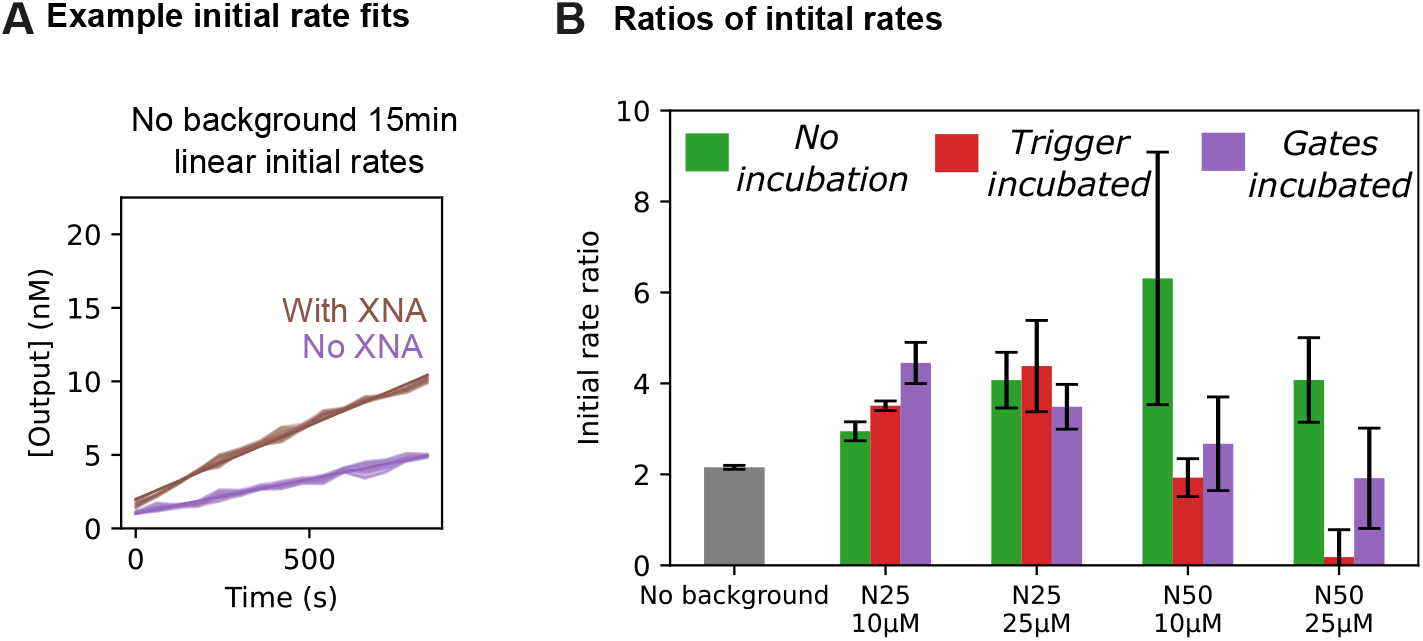
Alternative analysis of the kinetic effect caused by incorporation of the non-natural isoC:isoG base pair. (A) Example of a linear initial rate fit to the collected fluorescence data over the time period of 15 minutes. (B) Ratios of the initial rates across the tested conditions. The ratio is calculated by dividing the rate for the case with non-natural bases by the rate for the case without non-natural bases. The trend for the initial rate ration across random background length is qualitatively different than the bulk second-order rates from Figure 7. The large error in the N50 10*µ*M case stems from a very small slope magnitude in the natural case compounding in a large propegated error in the ratio of initial rates.

**Fig. S4.**
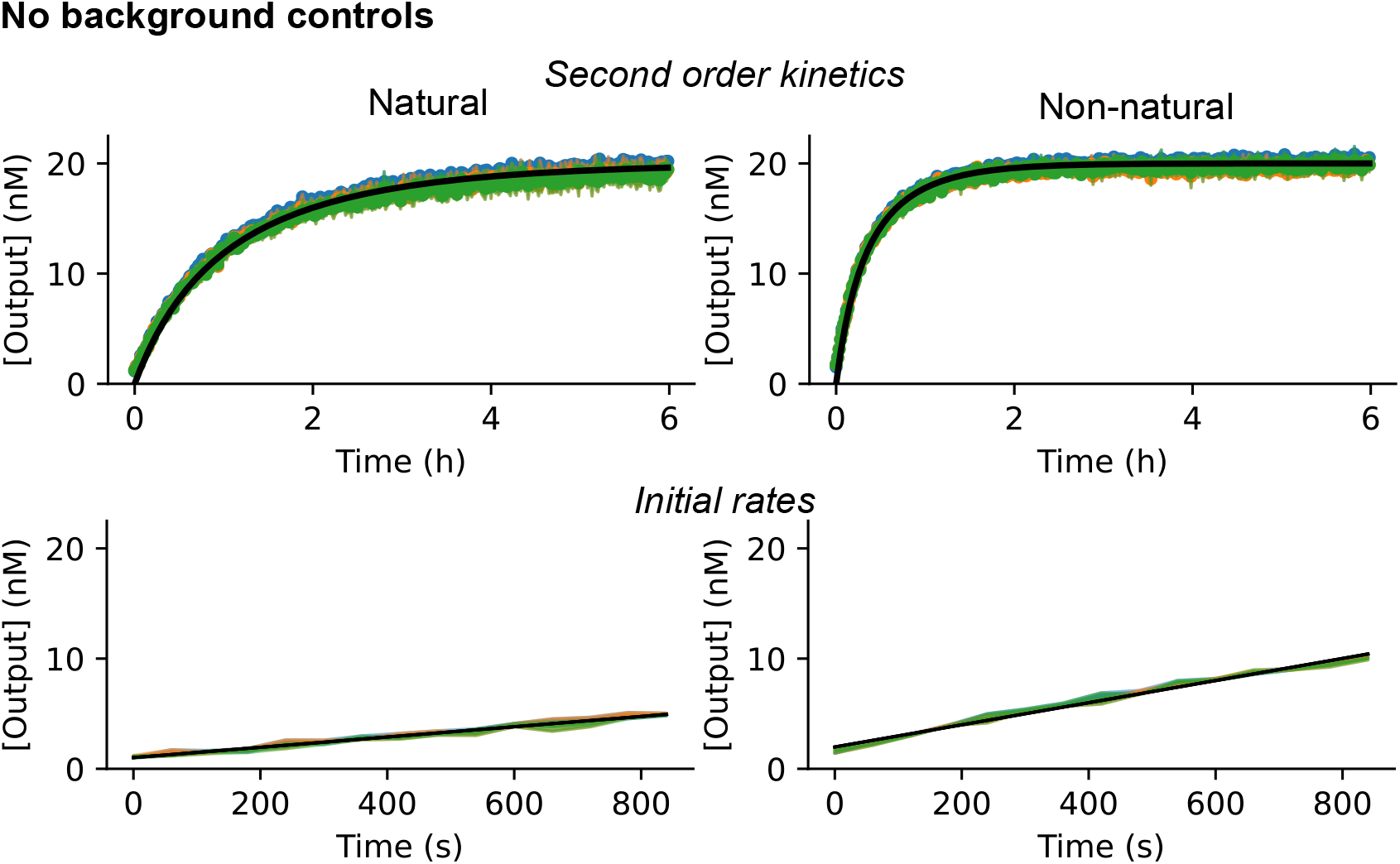
Kinetic curves for control data (without background) for the 5’ toehold system fit with an effective second-order rate constant (top) or linear initial rate (bottom). Data is presented for the natural control in the left column and the non-natural system in the right column. The three different colored curves represent the three sets of averaged duplicates used to estimate the variance of fit parameters by the jackknife method. The black curves represent the second-order or initial rate models plotted with the average parameter for the fit determined through the jackknife method.

**Fig. S5.**
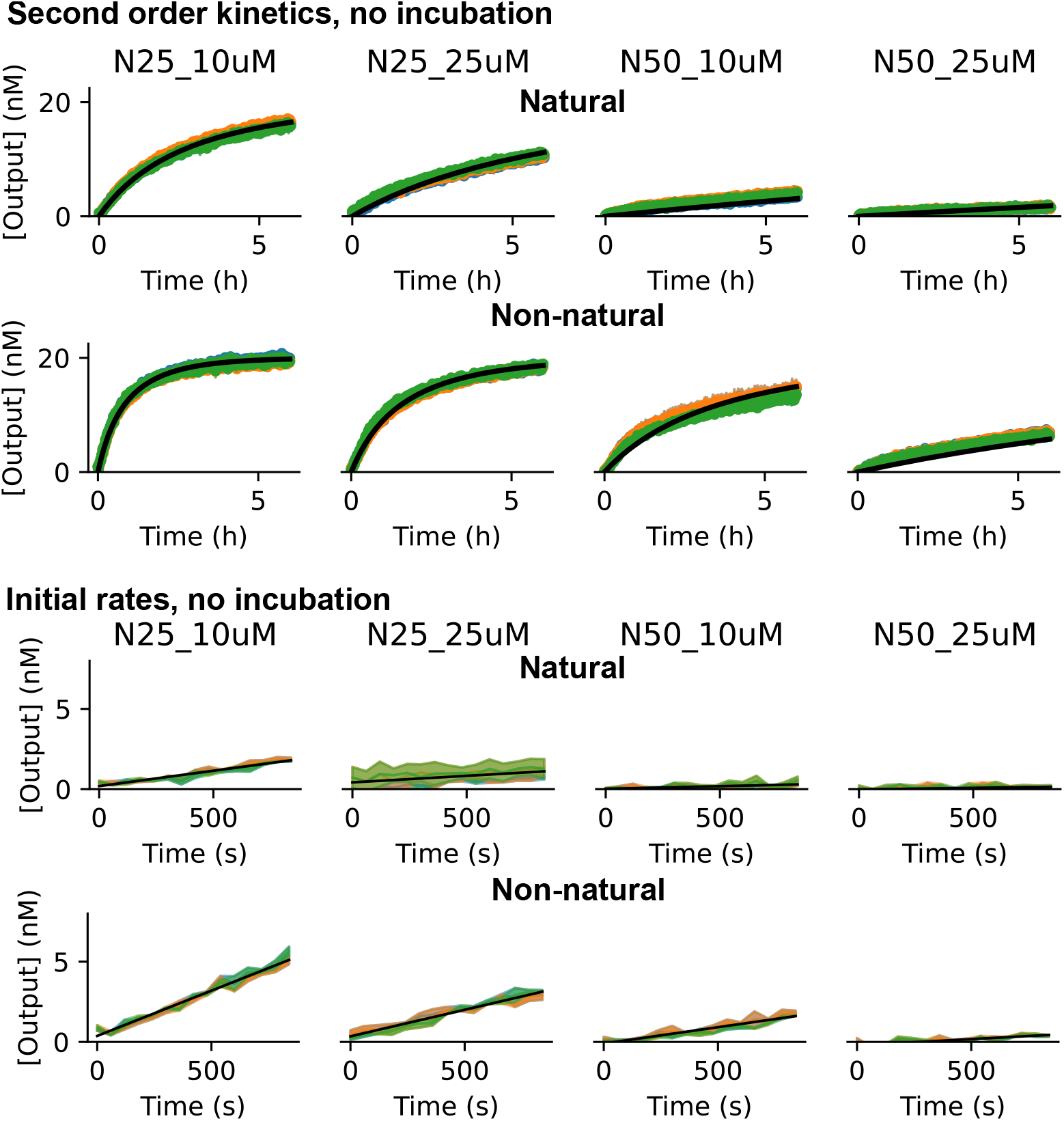
Kinetic curves for the 5’ toehold system added without incubation to different concentrations and lengths of background and fit with an effective second-order rate constant (top two rows) or linear initial rate (bottom two rows). Data is presented for the natural control in the first and third rows and the non-natural system in the second and fourth rows. The three different colored curves represent the three sets of averaged duplicates used to estimate the variance of fit parameters by the jackknife method. The black curves represent the second-order or initial rate models plotted with the average parameter for the fit determined through the jackknife method.

**Fig. S6.**
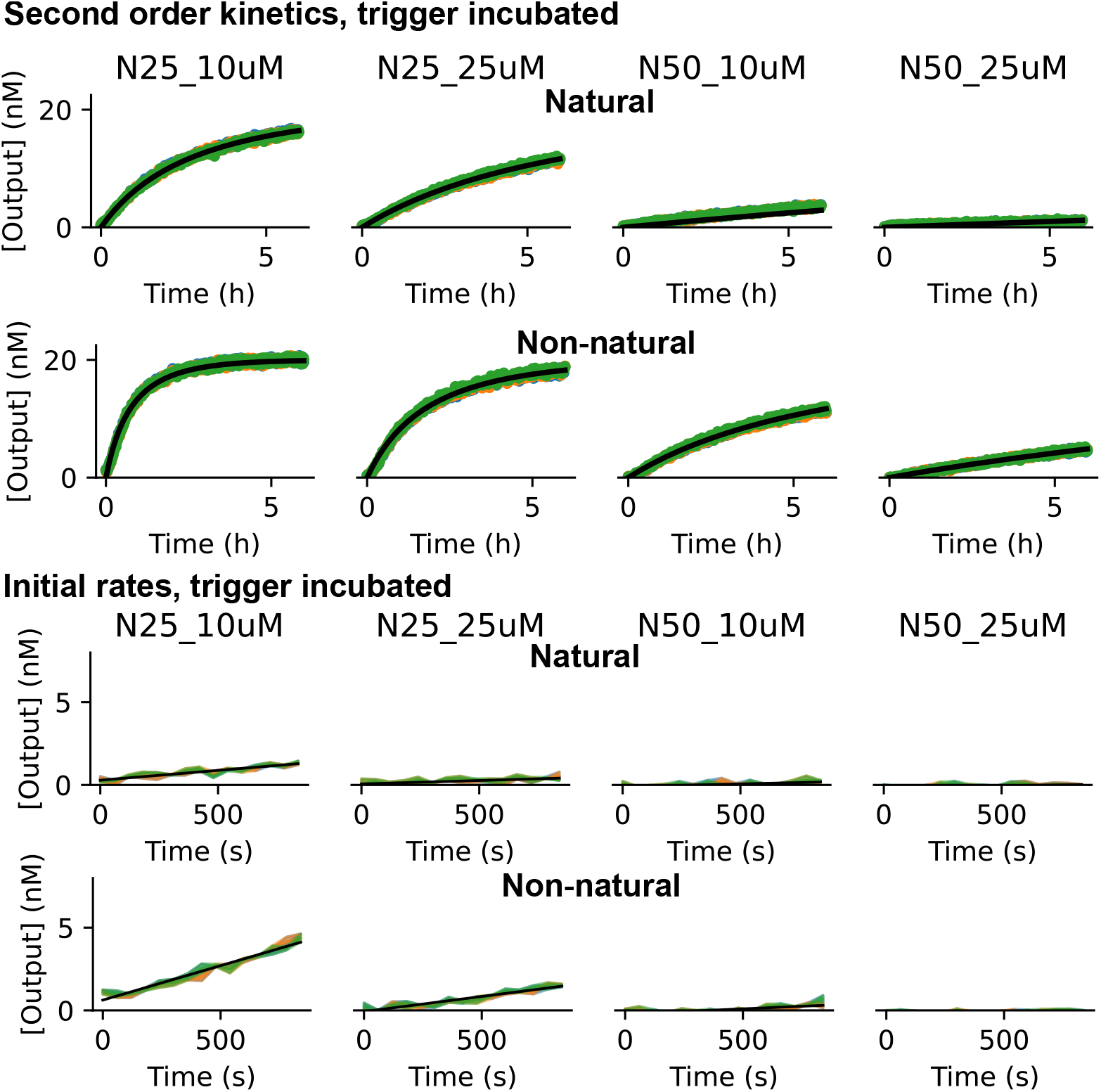
Kinetic curves for the 5’ toehold system with the trigger incubated in different concentrations and lengths of background and fit with an effective second-order rate constant (top two rows) or linear initial rate (bottom two rows). Data is presented for the natural control in the first and third rows and the non-natural system in the second and fourth rows. The three different colored curves represent the three sets of averaged duplicates used to estimate the variance of fit parameters by the jackknife method. The black curves represent the second-order or initial rate models plotted with the average parameter for the fit determined through the jackknife method.

**Fig. S7.**
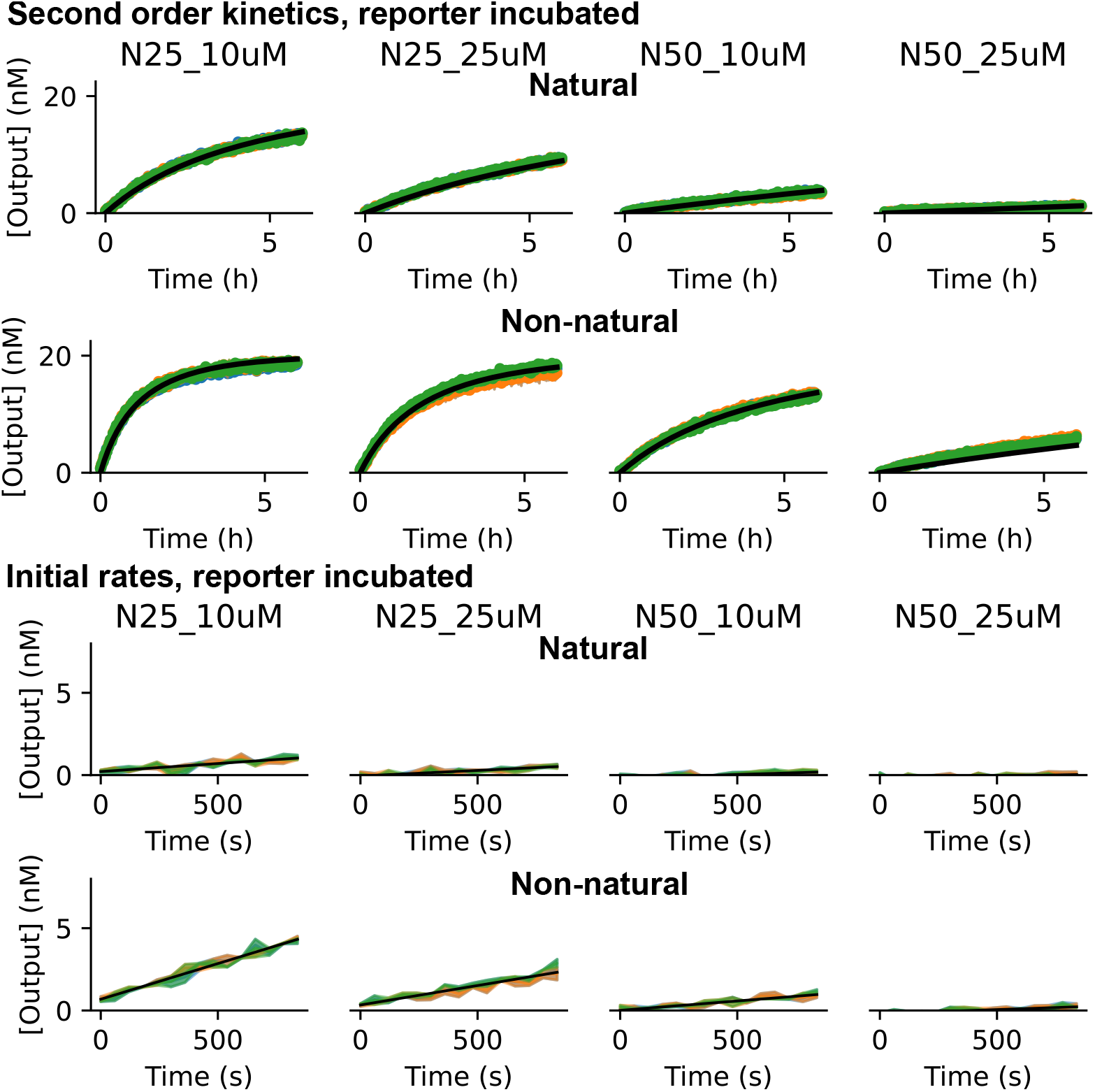
Kinetic curves for the 5’ toehold system with the reporter incubated in different concentrations and lengths of background and fit with an effective second-order rate constant (top two rows) or linear initial rate (bottom two rows). Data is presented for the natural control in the first and third rows and the non-natural system in the second and fourth rows. The three different colored curves represent the three sets of averaged duplicates used to estimate the variance of fit parameters by the jackknife method. The black curves represent the second-order or initial rate models plotted with the average parameter for the fit determined through the jackknife method.

**Table S2.**
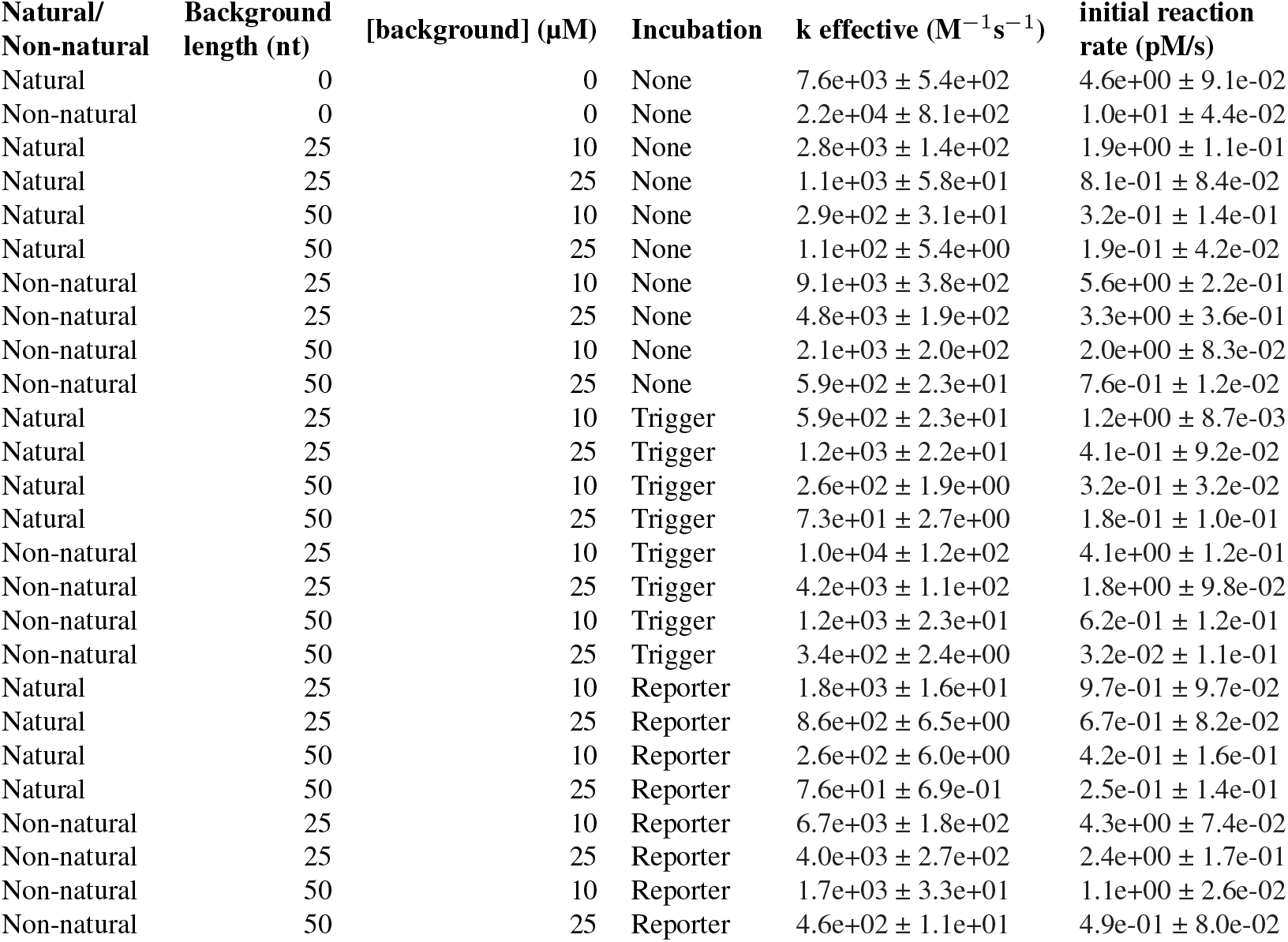
Effective rate constants and initial rates for each condition tested in the 5’ toehold system.

| Natural/<br>Non-natural | Background<br>length (nt) | [background] ($\mu\text{M}$) | Incubation | k effective ($\text{M}^{-1}\text{s}^{-1}$) | initial reaction<br>rate (pM/s) |
| --- | --- | --- | --- | --- | --- |
| Natural | 0 | 0 | None | $7.6\text{e}+03 \pm 5.4\text{e}+02$ | $4.6\text{e}+00 \pm 9.1\text{e}-02$ |
| Non-natural | 0 | 0 | None | $2.2\text{e}+04 \pm 8.1\text{e}+02$ | $1.0\text{e}+01 \pm 4.4\text{e}-02$ |
| Natural | 25 | 10 | None | $2.8\text{e}+03 \pm 1.4\text{e}+02$ | $1.9\text{e}+00 \pm 1.1\text{e}-01$ |
| Natural | 25 | 25 | None | $1.1\text{e}+03 \pm 5.8\text{e}+01$ | $8.1\text{e}-01 \pm 8.4\text{e}-02$ |
| Natural | 50 | 10 | None | $2.9\text{e}+02 \pm 3.1\text{e}+01$ | $3.2\text{e}-01 \pm 1.4\text{e}-01$ |
| Natural | 50 | 25 | None | $1.1\text{e}+02 \pm 5.4\text{e}+00$ | $1.9\text{e}-01 \pm 4.2\text{e}-02$ |
| Non-natural | 25 | 10 | None | $9.1\text{e}+03 \pm 3.8\text{e}+02$ | $5.6\text{e}+00 \pm 2.2\text{e}-01$ |
| Non-natural | 25 | 25 | None | $4.8\text{e}+03 \pm 1.9\text{e}+02$ | $3.3\text{e}+00 \pm 3.6\text{e}-01$ |
| Non-natural | 50 | 10 | None | $2.1\text{e}+03 \pm 2.0\text{e}+02$ | $2.0\text{e}+00 \pm 8.3\text{e}-02$ |
| Non-natural | 50 | 25 | None | $5.9\text{e}+02 \pm 2.3\text{e}+01$ | $7.6\text{e}-01 \pm 1.2\text{e}-02$ |
| Natural | 25 | 10 | Trigger | $5.9\text{e}+02 \pm 2.3\text{e}+01$ | $1.2\text{e}+00 \pm 8.7\text{e}-03$ |
| Natural | 25 | 25 | Trigger | $1.2\text{e}+03 \pm 2.2\text{e}+01$ | $4.1\text{e}-01 \pm 9.2\text{e}-02$ |
| Natural | 50 | 10 | Trigger | $2.6\text{e}+02 \pm 1.9\text{e}+00$ | $3.2\text{e}-01 \pm 3.2\text{e}-02$ |
| Natural | 50 | 25 | Trigger | $7.3\text{e}+01 \pm 2.7\text{e}+00$ | $1.8\text{e}-01 \pm 1.0\text{e}-01$ |
| Non-natural | 25 | 10 | Trigger | $1.0\text{e}+04 \pm 1.2\text{e}+02$ | $4.1\text{e}+00 \pm 1.2\text{e}-01$ |
| Non-natural | 25 | 25 | Trigger | $4.2\text{e}+03 \pm 1.1\text{e}+02$ | $1.8\text{e}+00 \pm 9.8\text{e}-02$ |
| Non-natural | 50 | 10 | Trigger | $1.2\text{e}+03 \pm 2.3\text{e}+01$ | $6.2\text{e}-01 \pm 1.2\text{e}-01$ |
| Non-natural | 50 | 25 | Trigger | $3.4\text{e}+02 \pm 2.4\text{e}+00$ | $3.2\text{e}-02 \pm 1.1\text{e}-01$ |
| Natural | 25 | 10 | Reporter | $1.8\text{e}+03 \pm 1.6\text{e}+01$ | $9.7\text{e}-01 \pm 9.7\text{e}-02$ |
| Natural | 25 | 25 | Reporter | $8.6\text{e}+02 \pm 6.5\text{e}+00$ | $6.7\text{e}-01 \pm 8.2\text{e}-02$ |
| Natural | 50 | 10 | Reporter | $2.6\text{e}+02 \pm 6.0\text{e}+00$ | $4.2\text{e}-01 \pm 1.6\text{e}-01$ |
| Natural | 50 | 25 | Reporter | $7.6\text{e}+01 \pm 6.9\text{e}-01$ | $2.5\text{e}-01 \pm 1.4\text{e}-01$ |
| Non-natural | 25 | 10 | Reporter | $6.7\text{e}+03 \pm 1.8\text{e}+02$ | $4.3\text{e}+00 \pm 7.4\text{e}-02$ |
| Non-natural | 25 | 25 | Reporter | $4.0\text{e}+03 \pm 2.7\text{e}+02$ | $2.4\text{e}+00 \pm 1.7\text{e}-01$ |
| Non-natural | 50 | 10 | Reporter | $1.7\text{e}+03 \pm 3.3\text{e}+01$ | $1.1\text{e}+00 \pm 2.6\text{e}-02$ |
| Non-natural | 50 | 25 | Reporter | $4.6\text{e}+02 \pm 1.1\text{e}+01$ | $4.9\text{e}-01 \pm 8.0\text{e}-02$ |

## Supplementary Note 3: Reporter rate constants, 3’ toehold system

We both analyzed the effect a non-natural base substitution in the toehold and the effect of a natural:non-natural mismatch on the rate constant of a single strand displacement reaction. We fit data from both reporters, as well as the non-natural reporter triggered with the natural trigger, to the following chemical reaction network (CRN) model:

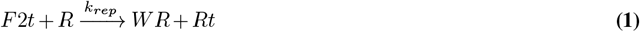

whose mass-action kinetics are defined by the following ordinary differential equation (ODE) model.

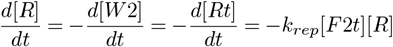

Firstly, the effective concentrations of each reporter was determined as described in Methods . Then the above ODE was fit to kinetic curves for the reporter triggered at three sub-stoichiometric concentrations to determine *k*_*rep*_. Since the reporter implements a strand displacement reaction rather than strand exchange, we approximate this to be an irreversible reaction. The experimental conditions and rate constant for each reporter analysis are presented in figure S8.

**Fig. S8.**
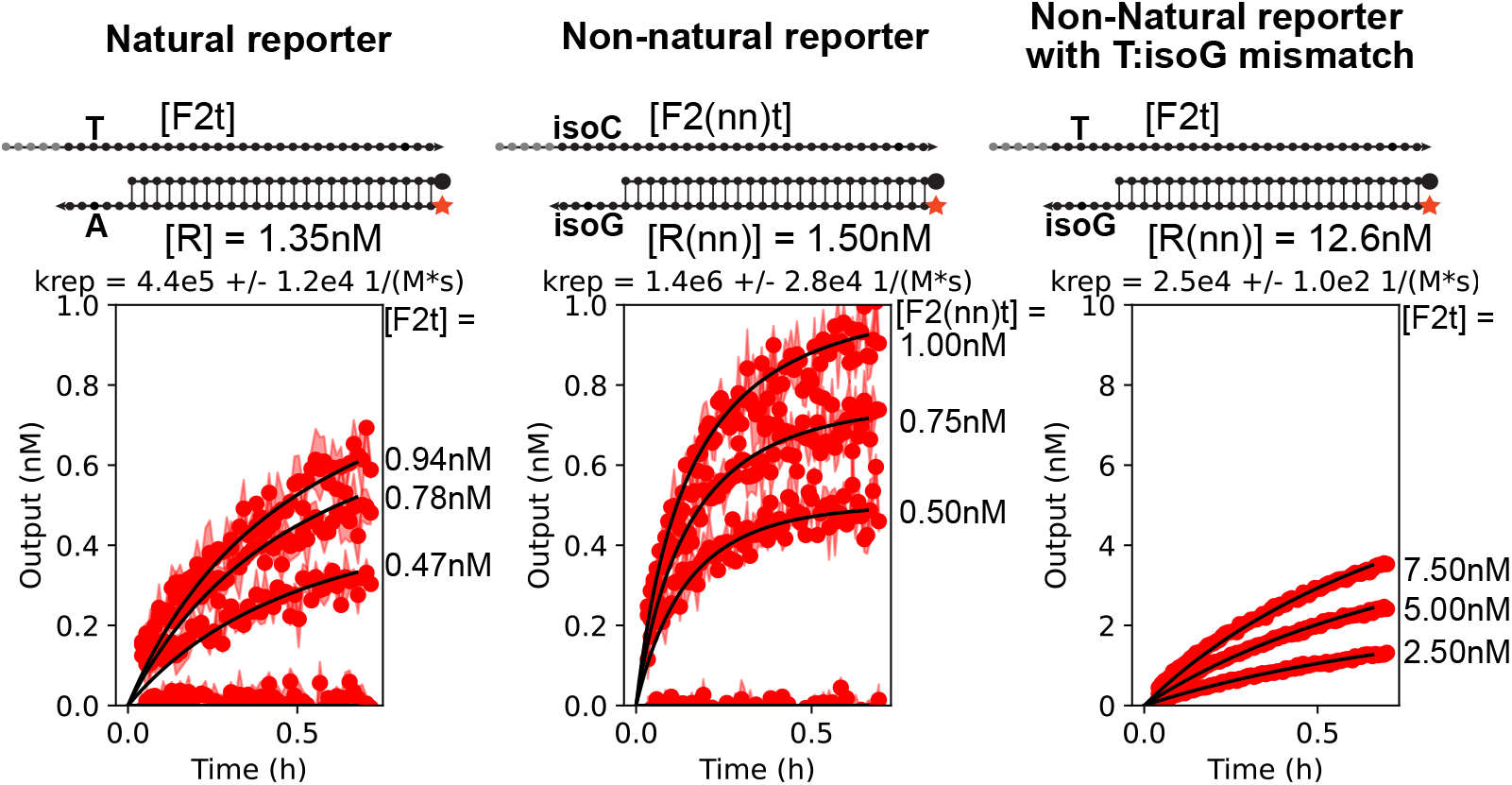
Rate constants for the 3’ toehold cascade’s natural reporter (left), non-natural reporter (center), and non-natural reporter triggered with natural trigger (right). The concentration of reporter and trigger used in fitting the model are shown for each system. Black curves indicate the model fits.

## Supplementary Note 4: Rate constants for each complex in both 3’ toehold cascades

To analyze the effect of XNA introduction on each gate in the cascade, we fit rate constants for each step of the natural and non-natural variants of the 5’ and 3’ toehold cascades. Firstly, as part of previous work, we fit both cascades to the following chemical reaction network (CRN) model:

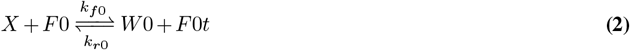

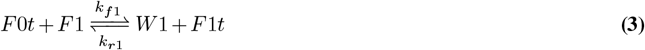

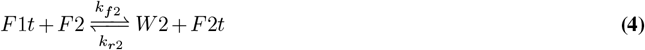

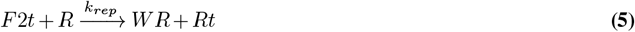

whose mass-action kinetics are defined by the following ordinary differential equation (ODE) model.

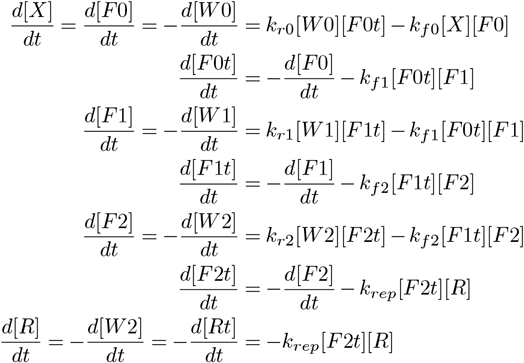

Firstly, the effective concentrations of each gate in the systems were determined as described in Methods, which alongside absorbance measurements of the trigger strands taken at 260nm using a Thermo Scientific™ NanoDrop™ One Microvolume UV-Vis Spectrophotometer were used determine the concentration of each species in the starting system for fitting each rate. Next the F0 and R gates were triggered with F1-t at three sub-stoichiometric concentrations. Using the established *k*_*rep*_ it was possible to fit a single parameter *k*_f2_, and use the detailed balance relationship ^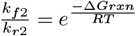^ where Δ*Grxn* is the free energy change of the reaction, R is the ideal gas constant, and T is the temperature– to determine *k*_*r*2_. We continues this procedure for the remaining two gates, analyzing each reaction gate one at a time and using the previously fit rate constants and detailed balance to reduce the problem to a single parameter fit at each step. Since we fit reverse rate constants of effectively 0 for the natural system, and the non-natural additions are expected to provide even more thermodynamic drive– increasing the ratio of the forward rate constant to the reverse rate constant, we assumed all reverse rate constants to be zero for the non-natural system. The fit-curves, rate constants, and experimental concentrations for each reaction are presented in figure S9. Since compare the new modeling for the non-natural cascade, to a previously collected model for the natural cascade, we chose to use the previously fit value of 4.1e5M^−1^s^−1^, which was collected using the same reporter strand synthesis and anneal as that used to collect the data to fit the subsequence rate constants, as opposed to the value of 4.1e5M^−1^s^−1^, which was collected at the time of this work using a new reporter synthesis and anneal. This is reasonable since different strand defects from synthesis and variations in strand stoichiometry can affect the rate constant.

**Fig. S9.**
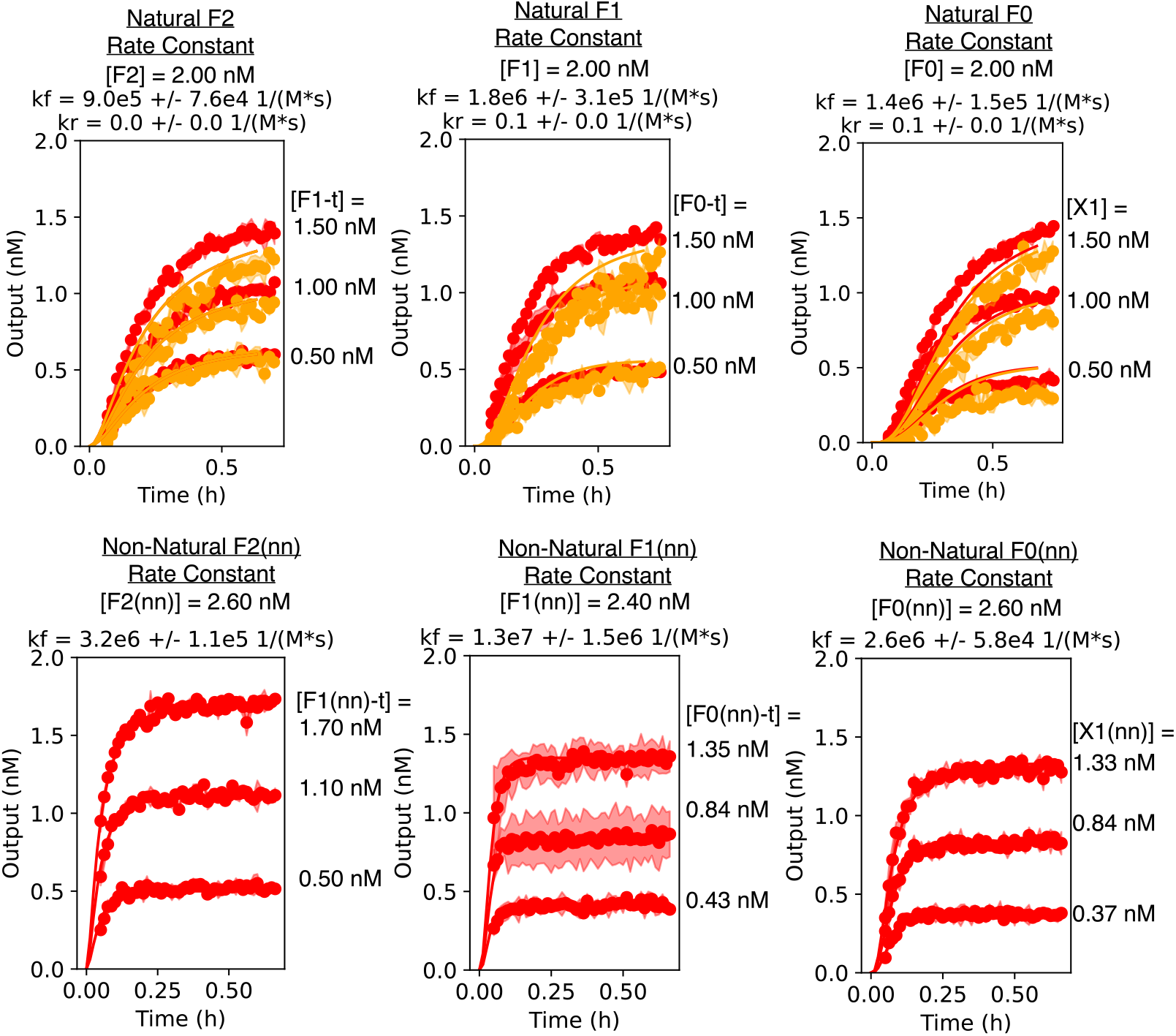
Rate constants for each complex in the natural 3’ toehold cascade (top row) and non-natural 3’ toehold cascade (bottom row). Each column shows the rate constant for a different complex in the cascade. The data for the natural cascade displays two different colors of curves, which correspond to technical replicated collected on different days as part of a prior study. Otherwise each curve represents a different concentration of trigger, with the trigger and concentration label. The concentration of gate of interest is labeled in the title of each plot. In determining the rate constant each gate, all upstream gates were omitted, the gate of interst was added at the concentration listed, and all downstream complexes were added at twice their listed concentration. The reporters were always added at 12nM for natural cascade and 17nM for the non-natural cascade.

## Supplementary Note 5: Tables of all sequences

**Table S3.**
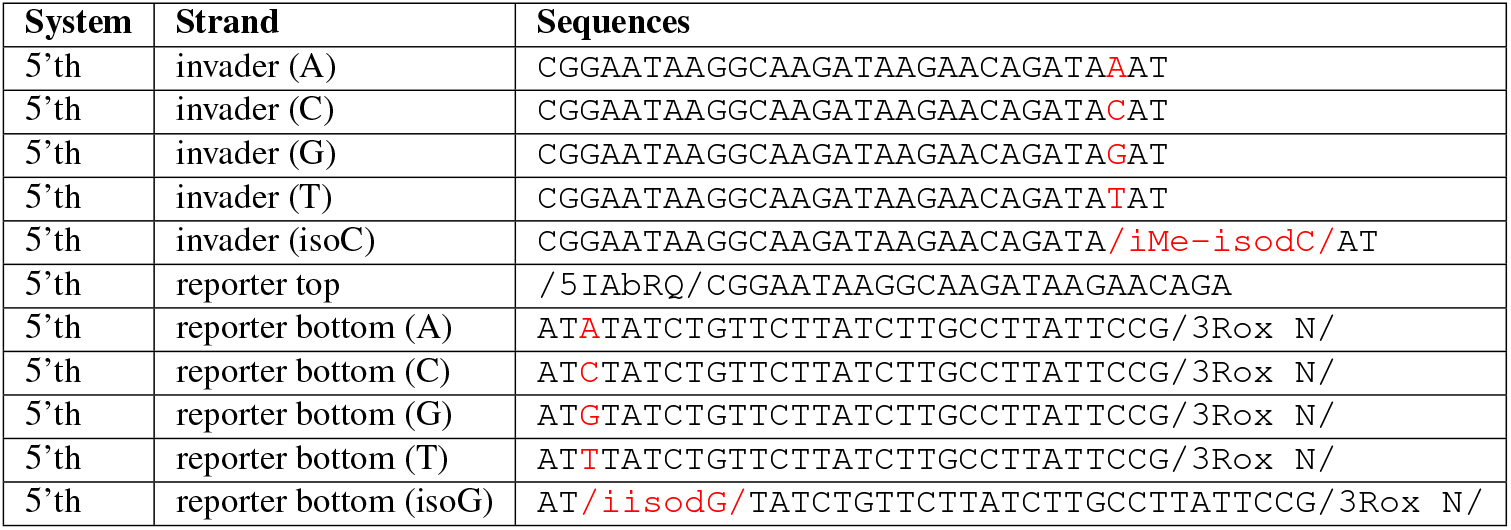
Promiscuity testing.

**Table S4.**
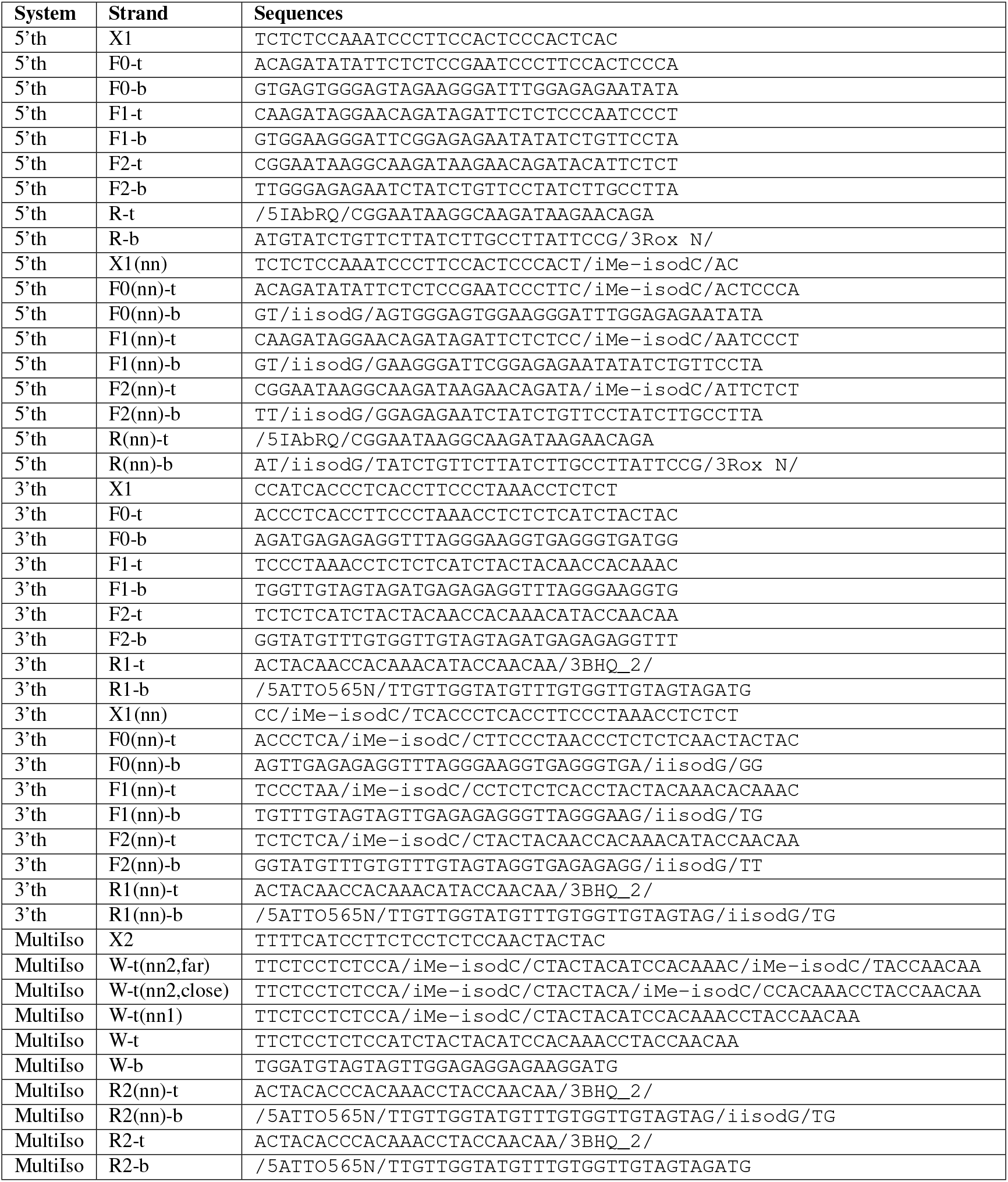
Displacement cascades.

